# The role of squid for food web structure and community-level metabolism

**DOI:** 10.1101/2023.07.14.549083

**Authors:** Rémy Denéchère, P. Daniël van Denderen, Ken H. Andersen

## Abstract

Squid differ from fish by their high growth rate, short life span and feeding behaviour. Their fast life strategy is thought to impose a high predation pressure on zooplankton, fish and other squid preys, and a rapid transfer of energy to upper trophic-levels of marine food webs. However, there is a lack of understanding of how squid’s fast life cycle affects the food-web structure, which is needed to project squid biomass across marine regions under shifting climatic conditions. Here, we examine the role of squid on community metabolism and biomass by collecting data on squid somatic growth and incorporating squid in a size- and trait-based fish community model. We show that squid have a 5 times higher average somatic growth rate than fish. Due to their high food demands, squid are constrained to regions of high pelagic secondary production. The presence of squid in these systems is associated with a reduction in total upper trophic level biomass. This decline is caused by an increase in community-level respiration losses associated with squid. Our results indicate that squid might have a large impact on ecosystem structure even at relatively low standing stock biomasses. Consequently, the recent proliferation of squid in ecosystems around the world is likely to have significant ecological and socio-economic impacts.

## Introduction

Squid populations have increased during the last six decades. This increase is thought to be due to the loss of top predators from fishing and rising temperatures (Doubleday et al., 2016; Pecl and Jackson, 2008). Squid are an important fisheries resource representing about 4% of the global marine landings (Hunsicker et al., 2010; Arkhipkin et al., 2015; Rodhouse, 2005). Locally, squid can be very important for fisheries, for instance they contribute up to 55% of the landings in the Patagonian Shelf and 30% in the Gulf of California. Squid also contribute indirectly to fisheries as a food resource for large predatory fish (Hunsicker et al., 2010) and are a major food resource for some whales (Garibaldi and Podestà, 2014).

Squid reach body sizes comparable to large teleost fish (hereafter termed fish) but differ from fish by their rapid growth, short lifespan, and semelparous reproduction strategy. For example, the flying jumbo squid can reach 140 cm in mantle length in less than two years (Goicochea-Vigo et al., 2019). These characteristics of squid imply that they must feed voraciously (Rodhouse et al., 1998, Chap. 13), which must be associated with high metabolic demand. Squid high feeding rates lead to high predation pressure on their prey, and potential top-down control of their prey, and rapid transfer of mass towards higher trophic levels (Rodhouse and Nigmatullin, 1996; Coll et al., 2013; Smale, 1996). The high metabolic demands and short life span make squid highly sensitive to inter-annual fluctuations in population biomass, potentially due to variation in prey availability and temperature (Boyle and Boletzky, 1996) and consequently high population variability.

Beside the physiological differences between fish and squid, there are also differences in the feeding niche. Both squid and fish increase their trophic niche as they grow in size. Squid have a diverse diet that shifts from small crustaceans to fish and other squid during ontogeny (Vovk, 1985; Phillips et al., 2003; MACY III, 1982). Squid inhabiting open oceans show a clear preference for mesopelagic fish (Hoving and Robison, 2016; Phillips et al., 2001), which indicates that they feed in deeper and darker waters than most pelagic fish predators. Predator-prey size ratios for squid are difficult to measure and estimates are not consistent among studies. For instance Vovk (1985) show that *Loligo palei* prefers prey of 4 to 24% of their length, whereas Hoving and Robison (2016) show that species of the genus *Gonatus* have a preference for prey of their own size. Nevertheless, squid differ from fish in their feeding mechanics. Fish swallow their prey whole, whereas squid use their long prehensile arms and beak to remove pieces of flesh from a prey. In this manner, squid can consume larger prey than fish. It is therefore likely that squid have a smaller prey:predator size ratio and, possibly, feed on a wider range of prey sizes than most fish.

The difference in life history and feeding niche between squid and fish implies that squid must play a different role than fish in structuring the ecosystem with potential consequences for top-down pressure, transfer of energy and biomass. Previous work on squid in food web models suggested the importance of squid in the marine food web (Morato et al., 2016). However, most models did not incorporate individual based processes of energy allocation with size (de la Chesnais et al., 2019; Coll et al., 2013), and therefore ignore linkages between the individual-, population-, and community-level dynamics (Persson et al., 2014; Arkhipkin, 2013). As a consequence, squid’s role in structuring ecosystems is not well understood.

In this study, we investigate how the distinct life history and feeding niche of squid, in comparison to fish, may impact population and community-level dynamics. To this end, we collected trait data on somatic growth rate *A* and adult:offspring size ratio *M*/*M*_0_ from several squid species, focusing on commercially important squid. We used this data to estimate two population metrics: the maximum population growth rate *r*_max_ and minimum sustainable resource level *R*^∗^. We compared both metrics to those previously derived for fish.To explore the community-level effects of squid on fish dynamics, we integrated squid in a size- and trait-based fish community model (FEISTY) (Petrik et al., 2019; van Denderen et al., 2021). This exploration was done for a range of modelled systems that varied in depth and ocean productivity. The structure and parameterization of the trait-based model reflected our aim of testing the community consequences resulting from the generic differences between squid and fish. The model and parameter values employed in this study do not aim to capture the entire ecological complexity and diversity of fish and squid species in natural systems. Rather, we represent the part of the community with the strongest impact on the community structure.

## Materials & Methods

We resolved the size structure of a squid population by developing a squid physiological model based on food-dependent growth and reproduction. We described all rates at the level of an individual squid as a function of its body mass *m* in units of wet weight. We used squid weight-at-age data to estimate the somatic growth rate and the maximum consumption rate. Assuming constant levels of food and predation, we derived the maximum population growth rate *r*_max_ and the minimum sustainable resource level *R*^∗^ (*m*) (Denéchère et al., 2022). We afterwards embedded squid into a size- and trait-based fish community model and explored the impact of squid on the fish community dynamics.

### Squid physiological model

We assume that squid physiology consists of encounter and consumption of food, losses from assimilation and standard metabolism, and allocation between growth and reproduction (Fig. 1A).

**Figure 1:**
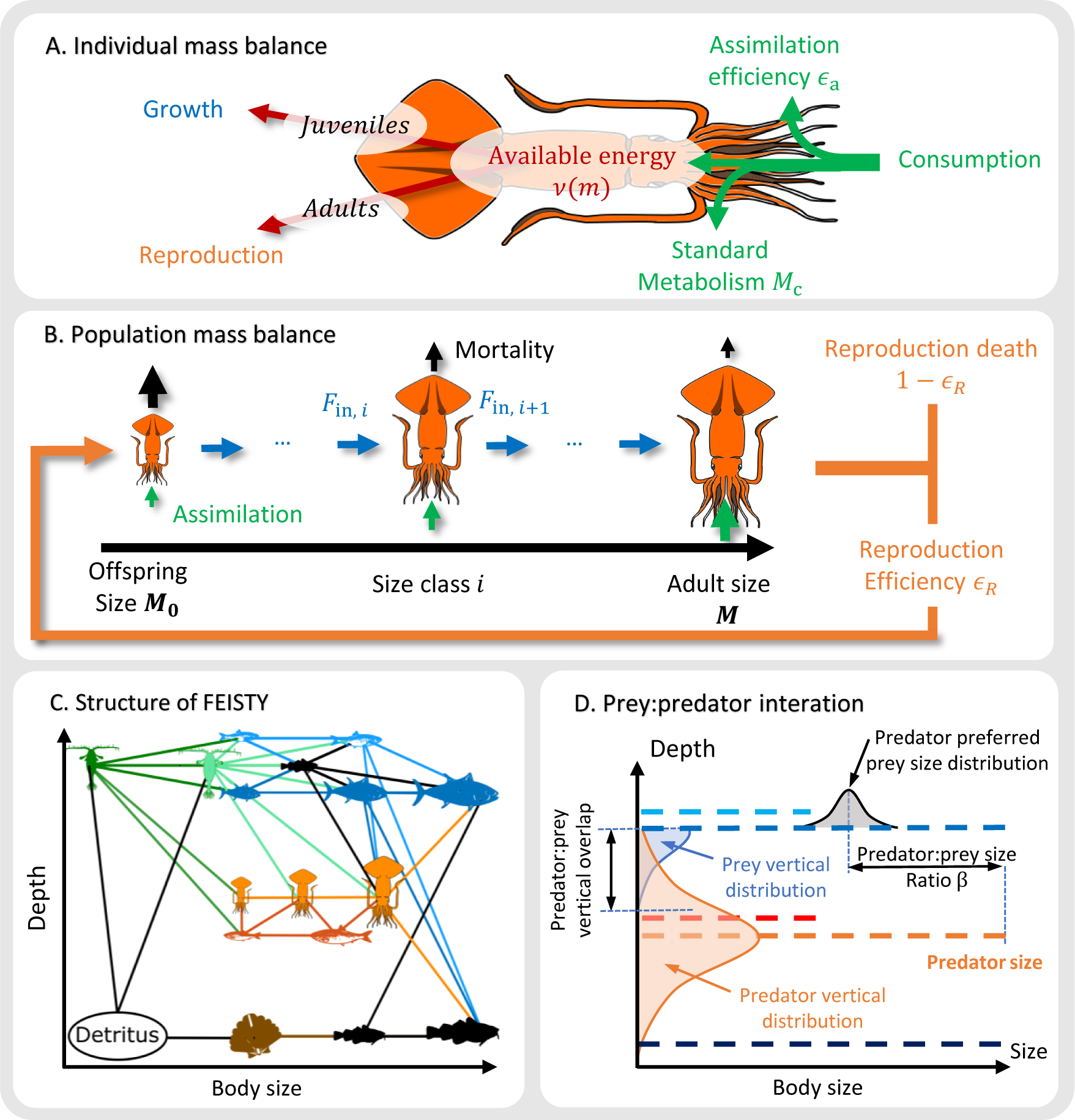
Representation of squid in the FEISTY model. (A) Individual physiological model of squid. (B) Population processes of squid. (C) Schematic representation of the size, depth and interaction of the functional groups in FEISTY. The model has three resources: small and large zooplankton (dark and light green respectively) and a benthic resource (brown); and five functional groups: small and large pelagic fish (light and dark blue respectively), demersal fish (black), mesopelagic fish (red) and squid (orange). (D) An example of vertical and size interaction in the FEISTY model.

**Figure 2:**
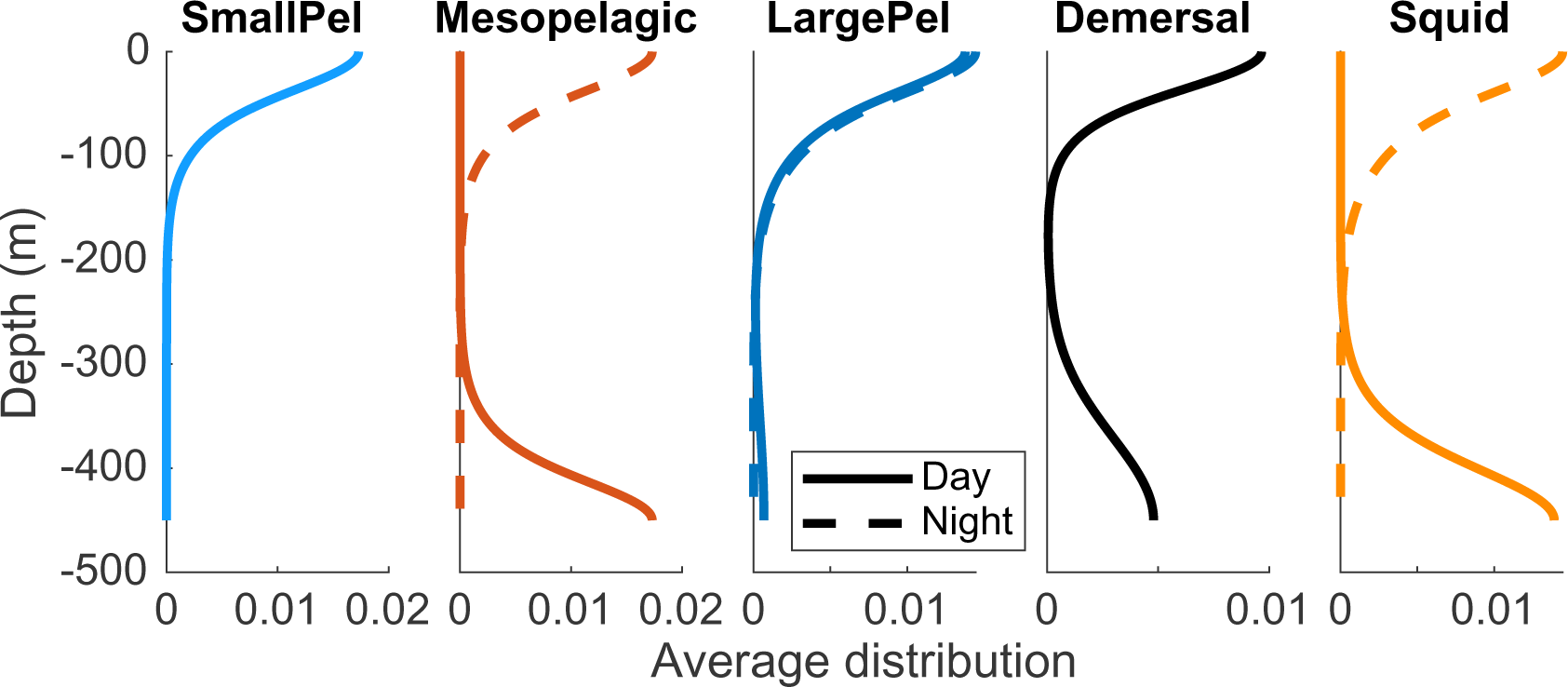
Day and night vertical distribution of fish and squid in the FEISTY framework. The vertical distribution is the average of all the life stages.

The encounter with food (mass per time) is the product of the search volume (volume per time) and the resource concentration *R* (mass per volume). We assumed that the search volume (area per mass per time) scales with body mass as *γm*^*q*^, with *γ* = 70 m^2^g^*q*^yr^−1^ the factor for search volume and *q* = 0.8 the exponent for search volume (Andersen and Beyer, 2006). The search volume is a proxy of the predation effort and represent a combination of swimming activity, predation mode (ambush or cruising), and the time spent feeding. Feeding is limited by the maximum consumption rate *hm*^*n*^ (mass per time), and the feeding level – the consumption relative to maximum consumption – is:

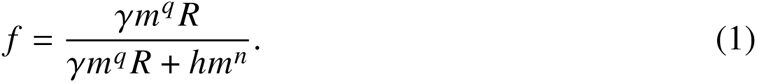

The consumed food *f hm*^*n*^ is assimilated with an efficiency *∈*_a_ = 0.7 and subjected to respiration losses. The available energy for growth and reproduction is then:

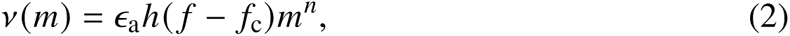

where respiration losses *∈*_a_*h f*_c_*m*^*n*^ are described as the “critical feeding levels” *f*_c_ ≈ 0.2 which is the respiration losses relative to the maximum assimilated consumption. Proportionality of metabolic cost with assimilation is assuming that metabolic cost scales with individual size with the same exponent that assimilation of energy (Denéchère et al., 2022). The value for *f*_c_ is estimated from fish (Andersen, 2019). The available energy is used for growth and reproduction. As squid are predominantly semelparous, juveniles use all available energy for somatic growth:

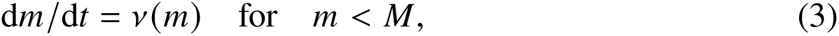

where *M* is the adult body mass. The reproductive rate of an adult is (no. offspring per time):

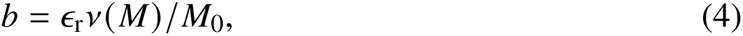

where the offspring size *M*_0_ is defined as the weight at hatching when offspring are independent of parental care (Neuheimer et al., 2015). The reproductive efficiency *∈*_r_ encompasses the additional costs of creating and storing eggs, egg mortality, and other costs of reproduction such as migration or forgone feeding. In the absence of information regarding *∈*_r_ for squid (Boyle and Rodhouse, 2008), notably the survival from egg release to hatching, we considered the reproduction efficiency equal for fish and squid, i.e., *∈*_r_ = 0.01 (Andersen, 2019).

Overall, the individual growth rate *A* and the short life cycle are the major differences between the life history parameters of fish and squid. Consequently, differences in the metabolic level and consumed food drive differences in growth and reproduction output.

### Estimating growth potential of squid

The coefficient for maximum consumption rate *h* represents the squid potential for somatic growth in our physiological model. Weighted it by the feeding level gives food-dependent growth. We collected weight-at-age data to calculate *h* for squid. This estimation requires us to assume an average constant feeding level *f* = *f*_0_ = 0.6. With this assumption somatic growth (2) can be rewritten:

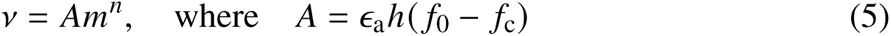

with *A* the somatic growth rate constant (units of mass^1−*n*^ per time) and *m* the mass of an individual. Classically, *A* is the growth coefficient in the von Bertalanffy growth type of model (see more detail for the link between *A* and our physiological model in Denéchère et al. (2022, Supplement D) or Andersen (2019, chap. 10). Solving (5), we find the mass as a function of age *t* (see Supplement A):

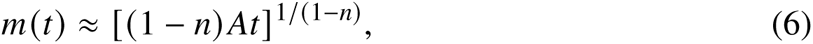

which we used to estimate the somatic growth rate *A* and then the maximum consumption coefficient *h* using eq. 5.

### Scaling from individuals to maximum population growth rate

We calculated two population level metrics to compare squid and fish: the maximum population growth rate *r*_max_ and the minimum sustainable resource level *R*^∗^. *r*_max_ describes the population growth rate in the absence of density dependence and can be interpreted as the invasive potential of a population. *R*^∗^ is the average resource level needed to sustain a population, i.e., when the density dependent population growth rate *r* = 0. *R*^∗^ represents the level of resource dependency or the level of intra-population density dependence.

Scaling from individual- to population-level metrics – *r*_max_ and *R*^∗^ – for squid followed the methodology from Denéchère et al. (2022). They derived *r*_max_ and *R*^∗^ for any size-structured population from individual-level metabolism and life history traits, i.e., mortality and growth. In the following we detailed the main assumptions and equations from Denéchère et al. (2022). We implemented *r*_max_ and *R*^∗^ with physiological parameters specific to both taxa (see parameters in Supplement A; Table. SA1).

*r*_max_ is estimated for a semelparous population assuming that the feeding level is constant *f* = *f*_0_ and that mortality is (units of per time) (Andersen and Beyer, 2006):

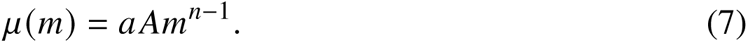

In this equation, mortality is proportional to the somatic growth coefficient *A* and assumes a linear trade-off between growth and mortality risk. This assumption accounts that faster growing species enhancing their foraging activities to fuel their grow and consequently increasing their own risk of predation, as is observed, e.g., among silversides (Lankford et al., 2001). The physiological level of the mortality *a* is a non-dimensional constant representing the ratio between mortality and mass-specific assimilated available energy *A*. We use *a* = 0.42 (Andersen, 2019). With this mortality, the maximum population growth rate is:

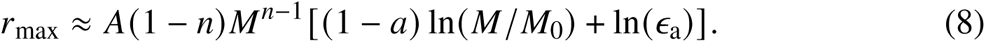

The approximation shows that the maximum population growth rate (per time) increases proportionally to the growth rate coefficient *A* and declines with increasing offspring size *M*_0_. We assumed that the resource environment *R* is constant and total mortality *μ*(*m*) only varies with size (but note that we include a full dynamic model where *R* and *μ* are dynamic for our food-web model).

*R*^∗^ is found by merging eq. 8 and 1 and solving for the resource level where *r*_max_ = 0:

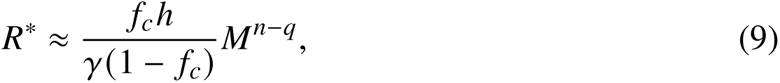

where *γ* and *q* are the parameters for search volume from eq. 1. We used the same *γ* and *q* values for squid and fish.

### Trait data collection

We collected data on three life history traits: somatic growth rate and the offspring and adult masses *M*_0_ and *M*. We estimated the somatic growth rate by fitting weight-at-age curves for 11 species of squid with a somatic growth model (see eq. 6 and references in Supplement A Table. SA1). The estimation of the somatic growth coefficient *A* did not show any relation with temperature (see Fig.SA3 & SA4) therefore we used the average *A* value of all species to implement our model. We obtained offspring and adult size from 27 squid species. Offspring size was defined as the weight at hatching when offspring are independent of parental care (Neuheimer et al., 2015). Adult size *M* was defined as the maximum recorded size for a species.

### Scaling to food-web structure

We used the squid physiological parameters to implement squid in the dynamic FEISTY framework (van Denderen et al., 2021). FEISTY describes how the food-web structure of a fish community changes across water depths and variations in secondary production (see Supplement B). FEISTY resolves several functional groups that differ in their adult *M* and offspring sizes *M*_0_, and vertical position in the water column. Feeding is defined by the size of individuals following eq. 1 and by the vertical habitat which determines the available resources *R* for each size class (Fig. 1C). Mortality is mainly due to predation emanating from the feeding of larger individuals from other functional groups or cannibalism. To represent the semelparity of squid, we added mortality after reproduction.

We included small and large pelagic fish, demersal fish, mesopelagic fish, and squid in the model. We describe below how squid were embedded (Fig. 1D). Further details about the fish functional groups can be found in van Denderen et al. (2021). Equation and parameters for fish are in Supplement B.

### Squid size-based population model

Both fish and squid populations are described by the biomass *B*_*i*_ (biomass per area g m^−2^) in different size classes, each characterised by the geometric mean body mass within the size class *m*_*i*_. Small fish species – small pelagic and mesopelagic – have 4 size classes and a maximum body mass of 250 gram. Large species – large pelagic, demersal and squid – have 6 size classes. Large pelagic and demersal fish have a maximum body size of 125 kg. As squid are generally smaller than large pelagic fish, we assumed that squid have an intermediate maximum size of 5.6 × 10^3^ g.

**Table 1:** Squid parameters used for *r*_max_ and FEISTY implementation.

| Symbol | Name | Value | Unit | Note |
| --- | --- | --- | --- | --- |
| $M$ | Adult weight | $5.6 \times 10^3$ | g | (1)&(2) |
| $M_0$ | Offspring weight | 0.01 | g | (1)&(2) |
| $n$ | Metabolic exponent | 2/3 | - | (see Supplement A) |
| $h$ | Coefficient for max. consumption rate | 100 | $\text{g}^n \text{yr}^{-1}$ | (3) |
| $\gamma$ | Factor for search volume | 70 | $\text{m}^2 \text{g}^q \text{yr}^{-1}$ | (3) |
| $q$ | Exponent for search volume | 0.8 | - | (3) |
| $f_c$ | Critical feeding level | 0.2 | - | (4) |
| $\beta$ | Predator:prey size ratio | 50 | - | Smaller than fish |
| $\sigma$ | Width of size preference | 1.3 | - | (3) |
| $\epsilon_a$ | Assimilation efficiency | 0.7 | - | (3) |
| $\epsilon_r$ | Reproduction efficiency | 0.01 | - | (3) |
| $a$ | Physiological level of mortality | 0.42 | - | For $r_{\max}$ |
|  |  | var. | - | For FEISTY |

Note: (1) Neuheimer et al. \(2015\); (2) Villanueva et al. \(2016\); (3) van Denderen et al. \(2021\); (4) Andersen and Beyer \(2006\)

The change in biomass *B*_*i*_ within a size class *i* are due to somatic growth in and out of the class, biomass accumulation within the class, and predation:

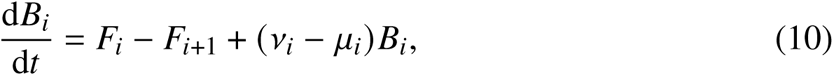

where *F*_*i*_ is the flux of biomass (biomass per time) from the lower size *m*_*i*−1_, and *F*_*i*+1_ is the flux of individuals growing into *i* + 1; *v*_*i*_ *B*_*i*_ is the accumulation of mass from feeding (Eq. 2), and *μ*_*i*_ *B*_*i*_ is the losses from predation.

The flux of mass between size classes is approximated by De Roos et al. (2008) as:

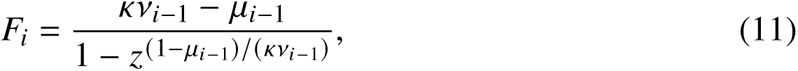

where *κ* describes the fraction of available energy invested in growth. As juvenile squid invests all energy in growth *κ* = 1. Adults invest all available energy in reproduction. Thus the flux out of the last size class *F*_x_ is used for reproduction, discounted by the reproductive efficiency eq. 4. The reproductive flux is routed into the first size class: *F*_1_ = *∈*_r_*F*_x_ assuming that adult squid die after reproduction (see Fig. 1B).

### Respiration cost

We assumed a trade-off between respiration loss *M*_c_ and consumption *h*, where *M*_c_ = *f*_c_ *hm*^*q*^ (2). As fish and squid have different somatic growth rates, and therefore respiration losses, we examined the effect of squid vs. fish abundance on community-level respiration. The respiration from a group *M*_tot_ is calculated as

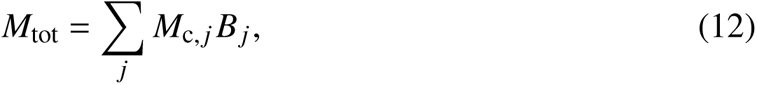

with *j* the size classes of a functional group

### Prey encounter

The squid population model is embedded in the food web (Fig. 1) via the resource *R*_*i*_ and predation mortality *μ*_*i*_of each squid size class *i*. The resource is determined by the available food from other groups (zooplankton, fish, and the squid population itself) and predation mortality is determined by the food consumption by all groups (fish through predation and squid through cannibalism).

Each group’s resource encounter is described as *R*_*i*_ = Σ*_j_ θ*_*i*,*j*_ *B*_*i*,*j*_, with *θ*_*i*,*j*_ the interaction matrix between squid of size *i* and a prey *j*, where *j* represent all size groups of all functional groups (including zooplankton and squid). The interaction matrix *θ*_*i*,*j*_ consists of two parts: the predator-prey size preference *θ*_size,*i*,*j*_ and the vertical overlap of each stage *θ*_*v*,*i*,*j*_, such that: *θ*_*i*,*j*_ = *θ*_size,*i*,*j*_ × *θ*_*v*,*i*,*j*_. The size preference is described by the optimal prey:predator size ratio *β* and the width of the size preference *σ* (Fig. 1C; Supplement B). Estimation of the size preference, the predator:prey size ratio *β* and width of the preference *σ*, is rarely conclusive for squid, though their preferred prey:predator size is larger than fish (Vovk, 1985; Hoving and Robison, 2016). We chose a *β* = 50 resulting in a preferred prey size 8 times larger than fish prey.

The vertical preference follows a bi-modal vertical distribution that represents diel vertical migration patterns of squid (Supplement B; Table. SB1). Oceanic squid dive into the twilight zone during the day and migrate to the surface at night (Roper and Young, 1975). For example, the jumbo squid (*Doscidicus gigas*) dives below 200 m depth during the day (William et al., 2006). The dive depth varies considerably between species and also, probably, depends on the resource availability. We therefore assumed that squid follow the day/night vertical movement of the migrating zooplankton and mesopelagic fish with a maximum concentration at similar depths. The vertical behaviour of coastal squid has received less attention, we assumed that squid in shelf waters also realise diel vertical migrations from the surface waters to the seafloor. All functional groups in FEISTY feed days and nights.

### Co-existence of squid in the FEISTY framework

Competitive exclusion between squid and fish groups is largely avoided by the differences in size and feeding niche between squid and fish. To avoid competitive exclusion between demersal fish and squid in shelf systems, we assumed that squid do not eat on benthic resources. This restriction was a necessary simplification of the empirical observations (see further discussion section).

### Food web analysis

We examined the equilibrium conditions of fish and squid for varying zooplankton productivity (g m^−2^ yr^−1^) and across sea floor depths (m). These variables also affect benthic resource productivity, which was changed accordingly (van Denderen et al., 2021) (Supplement B; Table. SB1 & SB2). We chose two depths, 50 and 2000 m (hereafter termed shelf and open ocean, respectively), to illustrate our results but the main results are robust when other depths were used. We varied the maximum zooplankton productivity for both small and large zooplankton from 10 to 150 gram wet weight per m-2 per year (for comparison the North Sea has an estimated productivity of 90 gram wet weight per m-2 per year for each zooplankton group that is available as prey for fish (Stock et al., 2017). For each depth and productivity combination, we ran the model for 150 years and averaged over the last 40 years (by which time the model had converged). Initial conditions of fish and squid biomass were 0.01 g for each size class and zooplankton and benthic biomass were 10% of the maximum resource carrying capacity used for the system. We verified that there were no alternative stable states by comparing simulation with increasing and decreasing zooplankton productivity.

Data and code are available on GitHub: https://github.com/RemyDenechere/The-role-of-squid-for-food-web-structure-and-community-level-metabolism/releases/tag/v0.beta

## Results

### Individual and population growth rate

Squid are fast growing species with average growth coefficient *A* = 27.9, compared to 5 g^1−*n*^ yr^−1^ for teleost fish (Fig. 3A). The egg mass of squid is independent of adult mass, just as it is for fish, but the average offspring mass is an order of magnitude higher, i.e., 0.01 and g for squid and fish respectively (Fig. 3B). The larger offspring size lowers maximum population growth rate, but their faster somatic growth rate more than compensates and squid have a higher maximum population growth rate than fish (Fig. 3C). The minimum sustainable resource *R*^∗^ is higher for squid than for fish suggesting that squid require higher food density to sustain their population size (Fig. 3D).

**Figure 3:**
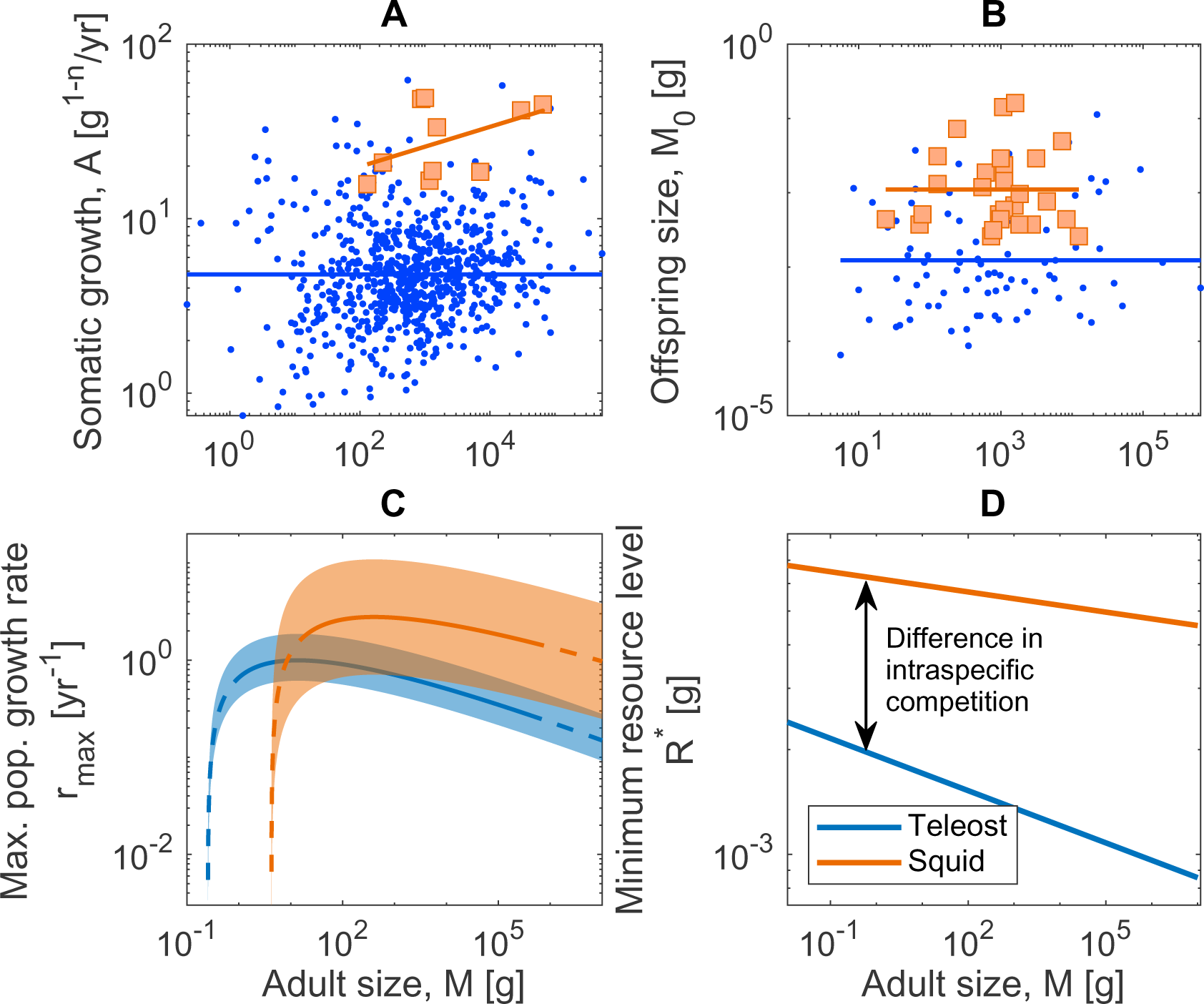
(A) Somatic growth rate *A*, (B) offspring weight *M*_0_, (C) maximum population growth rate *r*_max_, and (D) minimum resource level *R*^∗^ for squid and fish (orange and blue respectively) as a function of adult body size *M*.

### Food-web structure

Our results show that the presence or absence of squid is primarily driven by the productivity of zooplankton (Fig. 4 top panels). Squid occur in both shelf and open ocean when zooplankton productivity exceed 25 g m^2^ yr^−1^. The presence of squid leads to a decline in fish biomass and a decline in total community biomass.

**Figure 4:**
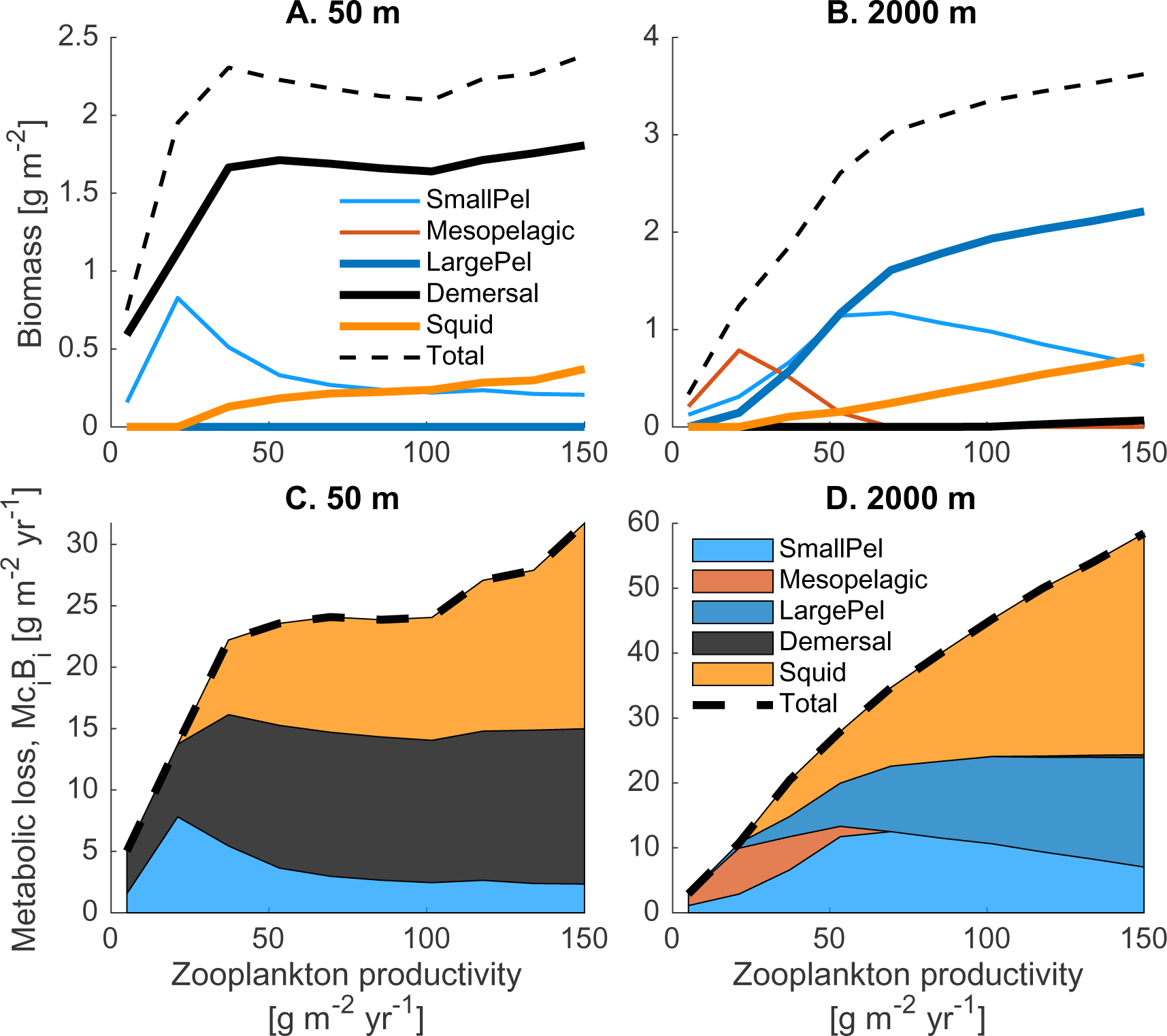
Biomass (top panels) and rate of metabolic loss (lower panels) of each functional group with increasing zooplankton productivity in the FEISTY framework for a shelf region (50 m depth) and an open ocean (2000 m depth) system. Thickness of the lines refers to the asymptotic size of the functional group. “Pel” in the legend refers to “pelagic”. Total metabolic cost per group is estimated as the sum of the metabolic cost per size

Both fish and squid switch trophic niche as they grow larger (Fig. 5). Squid and large pelagic fish shift from a zooplankton to a fish and squid diet, while small pelagic and mesopelagic fish shift from small to large zooplankton. Demersal fish switch from zooplankton to benthic prey and subsequently to fish and squid. As expected, demersal fish are dominant in shelf waters where they feed on both pelagic and benthic prey resources. In open ocean systems, mesopelagic fish are most dominant at relatively low zooplankton productivity and large pelagic fish at higher productivity (Fig. 4 top panels).

**Figure 5:**
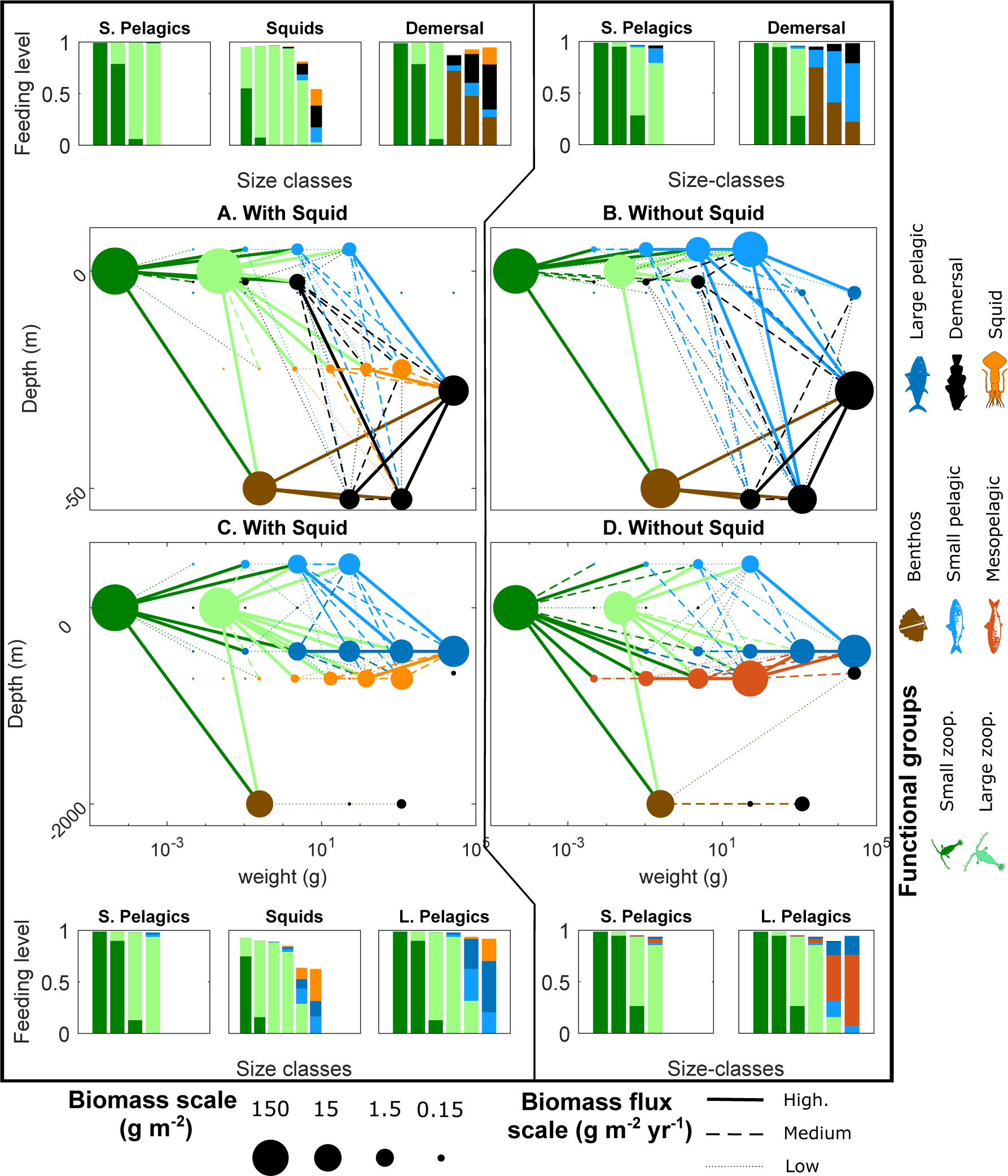
Comparison of the structure of the food web and feeding levels for systems with and without squid. Feeding levels of small pelagic, demersal and squid are presented for a shelf system (50 m depth) and small pelagic, large pelagic and squid for an open ocean system (2000 m depth). All the simulations are made for a zooplankton production of 130 g m^−2^ yr^−1^. For the sake of simplicity we only represent the 75 highest fluxes in each food web plot.

Squid biomass is relatively low in both shelf and open ocean up to the highest zoo-plankton productivity. Squid primarily feed on zooplankton and small pelagic fish in shelf areas (Fig. 5A; Squids) and on small pelagic and mesopelagic fish in the open ocean areas. Their feeding on mesopelagic fish results in a strong reduction of mesopelagic biomass compared to a system without squid (Fig. 5C & D).

Squid have low feeding levels, particularly as adults (Fig. 5A & C; Squids). Low feeding levels imply that a large fraction of their consumed prey is used to energetically cover the high respiration costs. All fish groups have higher feeding levels. This difference between squid and fish in feeding level shows that squid abundance is more resource limited (Fig. 5A & C; Squids) and fish abundance is more controlled by predators.

Squid is the biggest source of respiration losses among high trophic levels in the shelf system and a substantial fraction in the open ocean (Fig. 4 C and D), at least at high productivities. For instance, in a productive shelf system (50 m depth and 100 g m^−2^ yr^−1^ zooplankton productivity) squid account for 44 % of the total metabolic cost but only represent 9.1 % of the total biomass (Fig.4A and C). Similarly, in a productive open ocean (2000 m depth and 150 g m^−2^ yr^−1^ zooplankton productivity) squid account for 36.5 % of the total metabolic cost and 6.4 % of the total biomass (Fig.4B and D). Squid metabolic costs are higher than for fish due to their higher growth rate and since we assumed proportionality between standard metabolism *M*_c_ and individual growth rate *A* (2).

## Discussion

The squid species examined grow on average 5 times faster than fish (average somatic growth rates of 27.9 vs. 4.7 g^1−*n*^). Their faster growth leads to higher maximum population growth rate than fish, but also higher resource demand which constrains them to relatively productive systems. The presence of squid our a trait-based community model is associated with a reduction in total biomass in shelf systems and a reduction in the relative abundance of fish groups in open oceans. The reduction of fish biomass is not related to a drop in the feeding level of fish, suggesting that the effect of squid on fish is predominantly the result of predation by squid on fish and less due to competition for a shared resource. The model results further show that high respiration losses of squid decrease the upper total community biomass in systems where squid are present. These results indicate that, within the scope of squid species and feeding strategies examined in this study, squid play a significant role in shaping the ecosystem towards a state characterised by relatively low community biomass but high turnover of biomass.

### Differences in ecological success between fish and squid

The most conspicuous difference between fish and squid is in the somatic growth rate. Average squid growth is about 5 times faster than average fish growth and only the fastest growing fish mahi mahi (*Coryphaena hippurus*), have similar growth rates. What makes it possible for squid to grow so fast? It seems that their very active metabolism associated with high respiration from both cutaneous and gill respiration can sustain rapid growth (O’dor and Webber, 1986). Squid’s fast growth lead to high maximum population growth rates, which should allow them to out-compete slower growing fish populations. Yet, fish are more dominant than squid (Morato et al., 2016; Hunsicker et al., 2010). Fast growing species are also experiencing high risk of predation due to a more active lifestyle and risky behaviour resulting from their high food demand, which is thought to result in a dominance of species with “submaximal” population growth rate (Schramski et al., 2015). However, the trade-off between growth and predation risk is already accounted for in our calculation of maximum population growth rate so the higher predation risk is insufficient to explain the proliferation of fish over squid. We highlight that the fast living strategy of squid leads to a higher minimal resource requirement *R*^∗^ to maintain the population size (Fig. 3D). Consequently, squid populations are subjected to higher intra-specific competition and hence lower carrying capacity. We hypothesize that the reason for the lower success of squid compared to fish rests on their high resource requirements (high *R*^∗^).

### Dependency on zooplankton productivity

Our results predict that fast-growing squid should mainly inhabit productive regions as a result of their high demand in resource. This result is in agreement with some observations. For example, Boyle and Rodhouse (2008) argued that the most abundant Ommastrephidae squid inhabits productive regions, e.g., up-welling systems. To further validate these results, we reviewed EcoPath models that include cephalopod biomass estimates in different regions varying in zooplankton productivity and depth (Supplement C; Fig.SC1). Conversely to our results, EcoPath models show no increase in the proportion of cephalopod biomass (relative to fish biomass) at higher zooplankton productivity. This finding is in agreement with another study that found no relationship between cephalopod landings and primary production (Hunsicker et al., 2010). We have so far no explanation why there is such a difference between our theoretical expectation and these observations and predictions. It is, however, worth noticing that these studies examined total cephalopod biomass or landings and cephalopods may exhibit a wide range of life history strategies that differ from the fast-growing squid examined here (Seibel, 2007).

### Low standing stock biomass of fast living organisms

In accordance with our results, the review of EcoPath models show that cephalopods have a relatively low biomass compared with pelagic fish (ranging between 5 to 10% in most regions Supplement C; Fig. SC1). The low biomass of squid populations is the result of their high growth rate that imposes a low carrying capacity. Similarly, their semelparous strategy implies lower accumulation of adult biomass as for large fish. However, despite the low standing stock biomass the productivity (production of biomass per standing stock biomass) can still be high because of their high somatic growth rates.

One EcoPath model of the Azores region showed a much higher proportion of cephalopod biomass (see Supplement C; Fig. SC1B and Morato et al. (2016)). We expect that the cephalopods in this deep sea ecosystem, with seafloor depths up to 6000 meter, exhibit slower growth and lower metabolism than the squid studied here, see further (Seibel, 2007).

### Somatic growth as a key trait

We showed that somatic growth is a key trait to make predictions at the population and the ecosystem level, changing the relative abundance of fast- and slow-growing strategies. Unfortunately, few works address the importance of somatic growth rate for population dynamic. Stawitz and Essington (2019) shows the important contribution of somatic growth to the biomass of fish population. Since their somatic growth rate is a realised somatic growth, i.e., not corrected for food consumption, it is difficult to differentiate the growth strategy from the environmental condition in resource such as food availability. One could expect ecosystems with a high secondary production to exhibit a higher abundance of fast-growing species, with less standing stock biomass but higher productivity. At the macro-ecological level, there is some evidence that fish grow, on average, faster in productive regions (van Denderen et al., 2020; Morais and Bellwood, 2018). Yet, the signal is not very strong. For cephalopods it has been shown that landings are not correlated to primary productivity (Hunsicker et al., 2010), suggesting that the abundance of fast-growing strategies is not related to the system’s productivity. However, trophic transfer from the bottom to the top of ecosystems is complex and varies between regions. It is further known that fisheries catches (i.e. fish productivity) does not match with primary production (Ryther, 1969). Stock et al. (2017) showed that this mismatch could be solved by considering benthopelagic coupling and variation in transfer efficiency between regions. This finding shows that a high primary production might not reflect the high productivity of squid resources (secondary producers and small fish).

### Model uncertainty

The prediction of squid biomass and productivity in our dynamic model of ecosystem food-web (FEISTY-squid) is strongly dependent on the individual growth rate *A* evaluated for squid. Our estimation for this parameter rests upon a low number of species (11 species), and a poor estimation of the temperature effect on metabolic rates. Our estimation of somatic growth rate does not depend on temperature. The species in the analysis are dominated by commercially important squid, which is probably biased towards highly productive fast growing species. The results of our study will differ in systems where the dominant squid species grow equally fast or slower than the dominant fish species. Thus, examining variation in individual growth rate *A* of squid and fish species across a range of marine systems may help to better parameterise the model.

Squids vertical strategy is well described for many open ocean species, which make diel vertical migrations and preferentially target mesopelagic fish (Roper and Young, 1975). Squid vertical behaviour in species living in shelf waters is less well described. Feeding data show that some squid species feed on benthic organisms at intermediate sizes, suggesting that some shelf squid could occupy the niche of demersal fish. Juveniles live closer to the sea bed and transition to more pelagic niche when reaching the adult size (Vovk, 1985). In our model analysis, we decided to ignore squid feeding on benthic organisms to allow for niche differentiation between demersal fish and squid. Allowing squid to feed on benthos on shelf systems in the model would result in competitive exclusion of demersal fish at high resource productivities (not shown). This indicates that there are differences in the feeding niche between squid and demersal fish that we do not describe adequately, such as a higher feeding efficiency or time spent feeding on benthic resources.

In our model analysis, we assumed that squid are strictly semelparous. However, several studies have shown that some squid species may exhibit several spawning events during their life-span (Pecl, 2000; Hernández-Muñoz et al., 2016; Pérez-Palafox et al., 2019). These studies are controversial since they are unable to show truly iteroparous strategies in squid with gonad regeneration. The squid species are instead releasing already formed eggs several times during the season (Laptikhovsky et al., 2019). Since model predictions are primarily driven by the observed growth differences between fish and squid, implementing an iteroparous strategy for squid would not affect the qualitative results of the model.

## Conclusion and future direction

We show that squid have a potentially large impact on ecosystem structure and function even at relatively low biomass. We show that their fast growth and semelparous reproduction strategy drive this impact on ecosystem structure. We anticipate that the recent proliferation of squid in ecosystems around the world has likely caused significant ecological and socio-economic impacts to fisheries resources. This work could provides a framework to understand the expanding presence of squid across various ecosystems in relation to climate change and fishing activities.

## Acknowledgements

This work was supported by the Villum Kann Rasmussen Foundation Centre for Ocean Life, by the Independent Danish Research Foundation project Future Oceans. PDvD was funded by the European Union’s Horizon 2020 research and innovation programme under the Marie Sklodowska-Curie grant agreement No 101024886. We thanks Jeremy S. Collie at the Graduate School of Oceanography for the discussion and comments on our work. We declare no conflicts of interest.

## Statement of Authorship

All authors designed the model methodology. RD performed the research (data collection, data analysis, and coding simulation). All authors contributed to the writing.

## Supplement A Estimation of somatic growth rate

Cephalopods are usually semelparous (Boyle and Rodhouse, 2008). Their life cycle has two distinct phases: growth and reproduction. They die shortly after reproduction. We assume that the growth curves would fit a von bertalanffy growth model with a coefficient *k* = 0 (no investment in reproduction) with the following growth in mass per time:

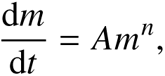

where *m* is the mass, *A* the growth coefficient and *n* the metabolic exponent. For the sake of simplicity we assume that *n* = 3/4. It can be rewritten:

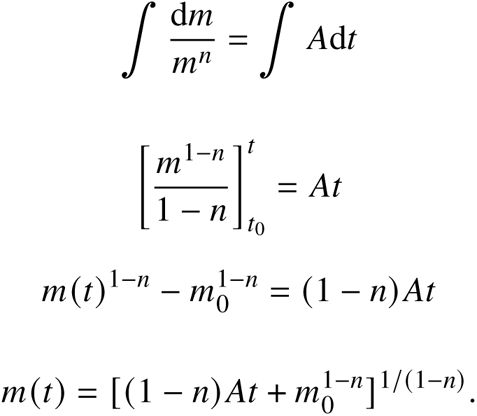

For the sake of simplicity we assume that offspring size *m*_0_ ≈ 0. The time dependent mass equation becomes:

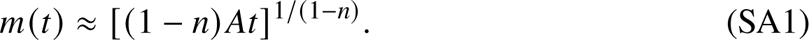

We collected weight and length at age curves (see Table. SA1) to estimate the growth rate of cephalopods. Most part of the individual growth data collected are expressed in either length or weight at age. To estimate individual somatic growth rate *A* we converted length at age curves in weight at age curves. We collected data of weight at length from (Kooijman, 2009) to estimate the conversion relationship from length to weight (Fig. SA1). We assumed that length *l* varies with weight *w* following: *w* = _c_ × *l*^*b*^. We found that length scales with weight with an exponent *b* = 2.2 suggesting that squid does not conserve the same volume structure throughout ontogeny.

Additionally, we estimated the somatic growth *A* for each temperature recorded with the data (Fig. SA3). The linear correlation between *A* and the temperature shows that the somatic growth rate is not scaling with temperature (p-value = 0.855; Fig SA4). Note that *A* is calculated on a sub-ensemble of individual for which the temperature is recorded. Therefore, the estimation of *A* might slightly differ from the one estimated in Fig. 3.

**Figure SA1:**
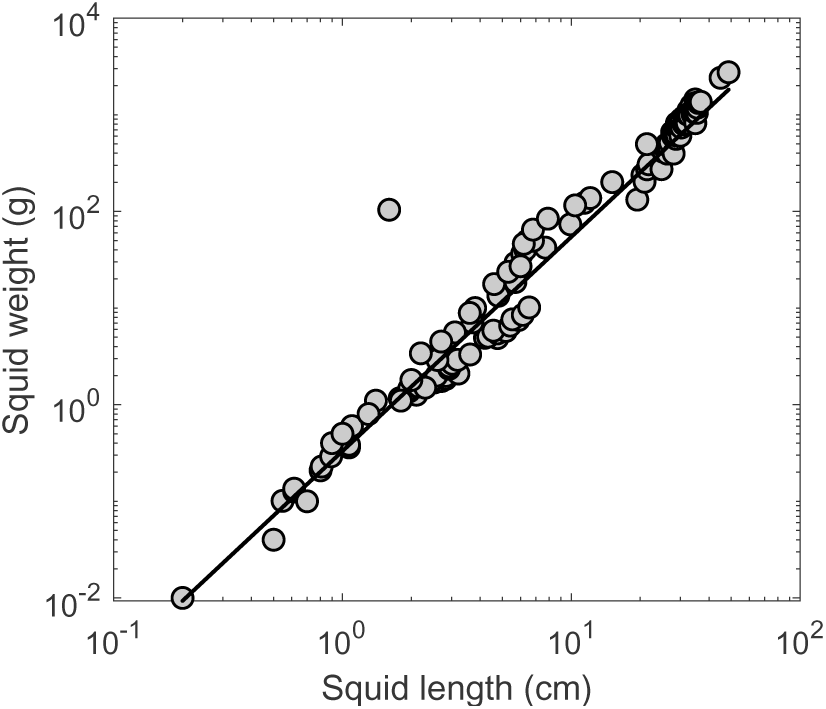
Relation between weight (g) and length (cm) for squid.

**Table SA1:** Summary of the data collected on somatic growth rate *A*, adult:offspring size ratio *M*/*M*_0_ and length at age coefficient.

| Symbol | Variable | Taxa | Average | Unit | Ref. |
| --- | --- | --- | --- | --- | --- |
| Offspring size | $M_0$ | squid | 0.01 | g | (1) & (2) |
|  |  | Teleost | 0.001 | g | (2) - (4) |
| Somatic growth rate | $A$ | squid | 27.9 | $\text{g}^{1-n} \text{y}^{-1}$ | (5) - (14) |
| | | Teleost | 5 | $\text{g}^{1-n} \text{y}^{-1}$ | (15) & (16) |
| Length-to-weight coefficient | $c$ | squid | 0.33 | - | (5) & (17) |
| Length-to-weight exponent | $b$ | squid | 2.2 | - | |

References: (1) Villanueva et al. (2016); (2) Neuheimer et al. (2015); (3) Denéchère et al. (2022); (4) Andersen (2019); (5) Kooijman (2009); (6) Summers (1971); (7) Pecl (2000); (8) Miyahara et al. (2006); (9) Rodhouse and Hatfield (1990); (10) Agus et al. (2018); (11) Arkhipkin and Silvanovich (1997); (12) Fang et al. (2016); (13) Jackson and Domeier (2003); (14) Rosa et al. (2013); (15) Hutchings et al. (2012); (16) Froese and Pauly (2018); (17) Nabhitabhata (1995)

**Figure SA2:**
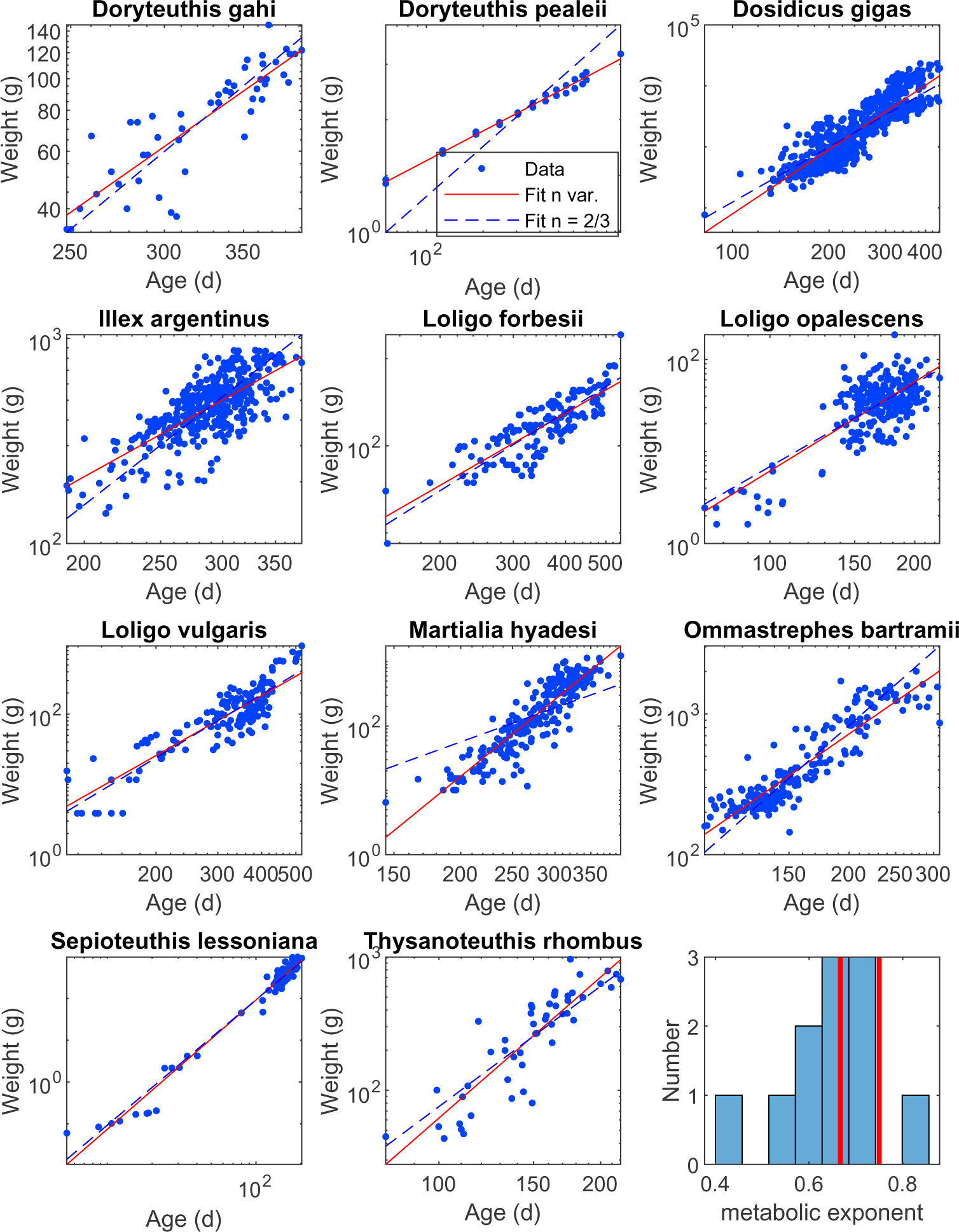
Weight at age data for 11 squid species. The red lines represents the linear regression with *n* the metabolic exponent as an estimated parameter. The blue dashed line represent the linear regression with fixed metabolic exponent: *n* = 2/3.

**Figure SA3:**
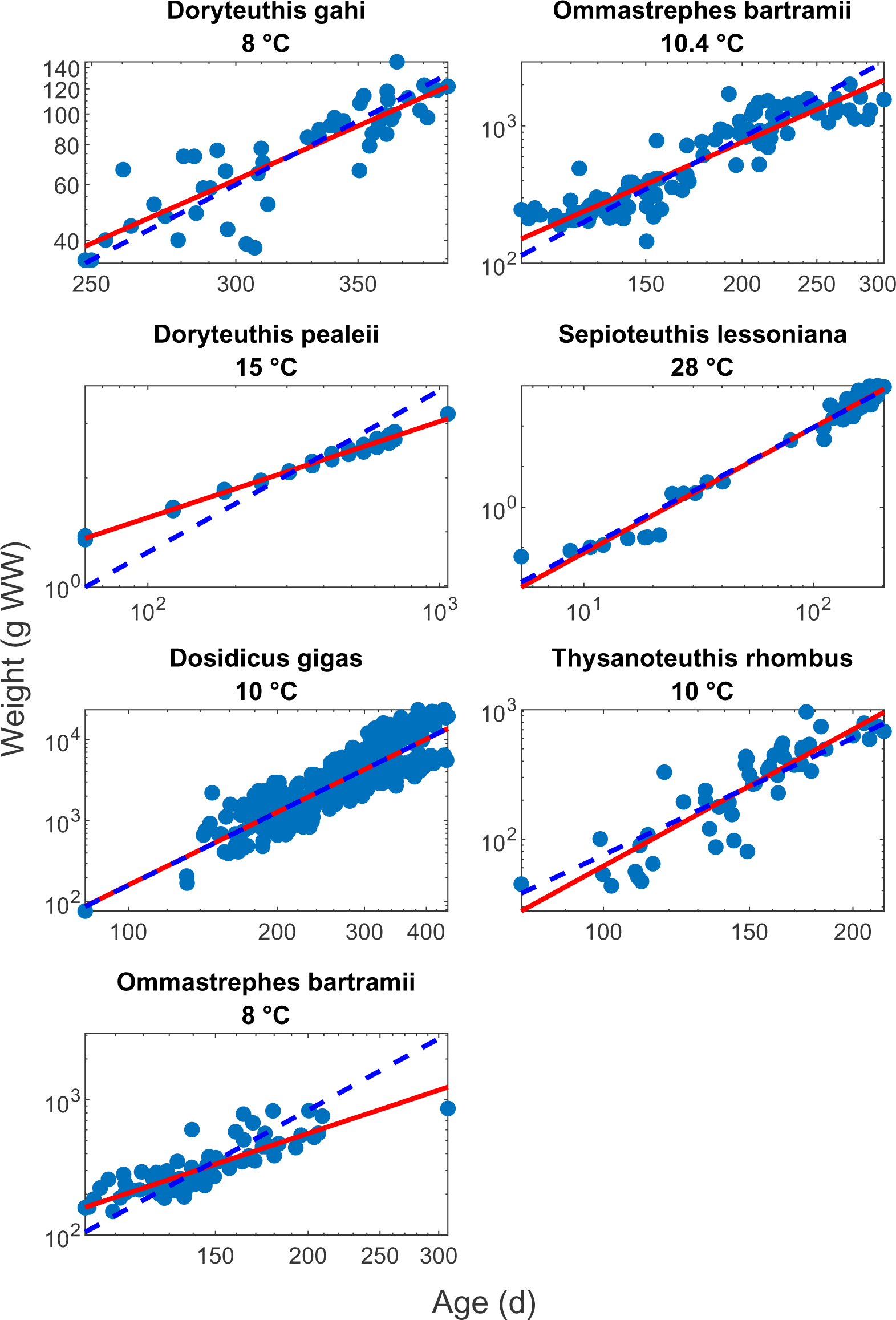
Weight at age for the species were temperature was reported. The red lines represents the linear regression with *n* the metabolic exponent as an estimated parameter. The blue dashed line represent the linear regression with fixed metabolic exponent: *n* = 2/3.

**Figure SA4:**
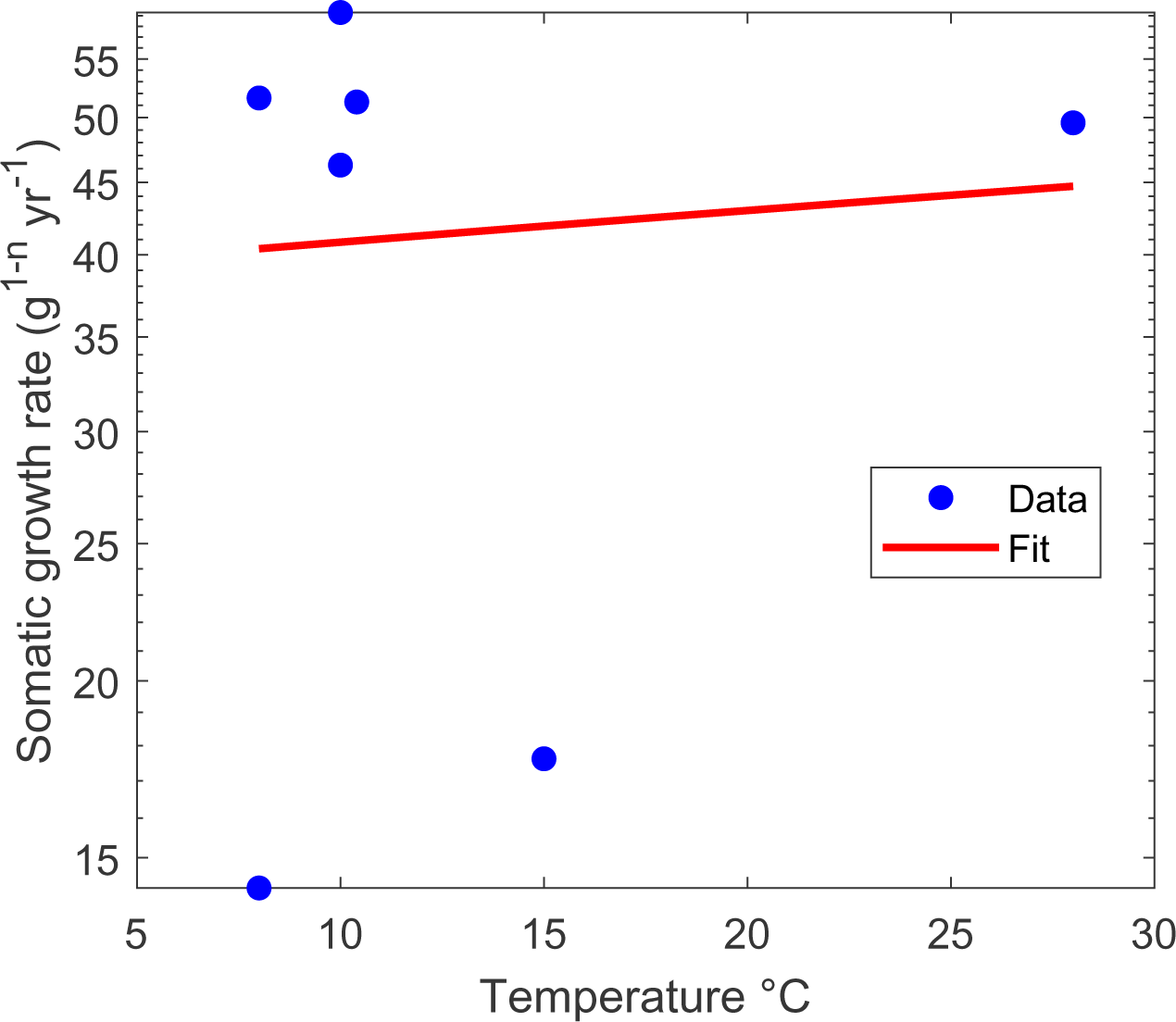
Estimation of somatic growth rate *A* (blue dots) from Fig.SA2 and eq. SA1. The linear regression (red line) does not show a significant relationship between the somatic growth rate and the temperature (p-value = 0.85).

## Supplement B Main equation of the FEISTY and parameters for the fish functional groups

**Table SB1:**
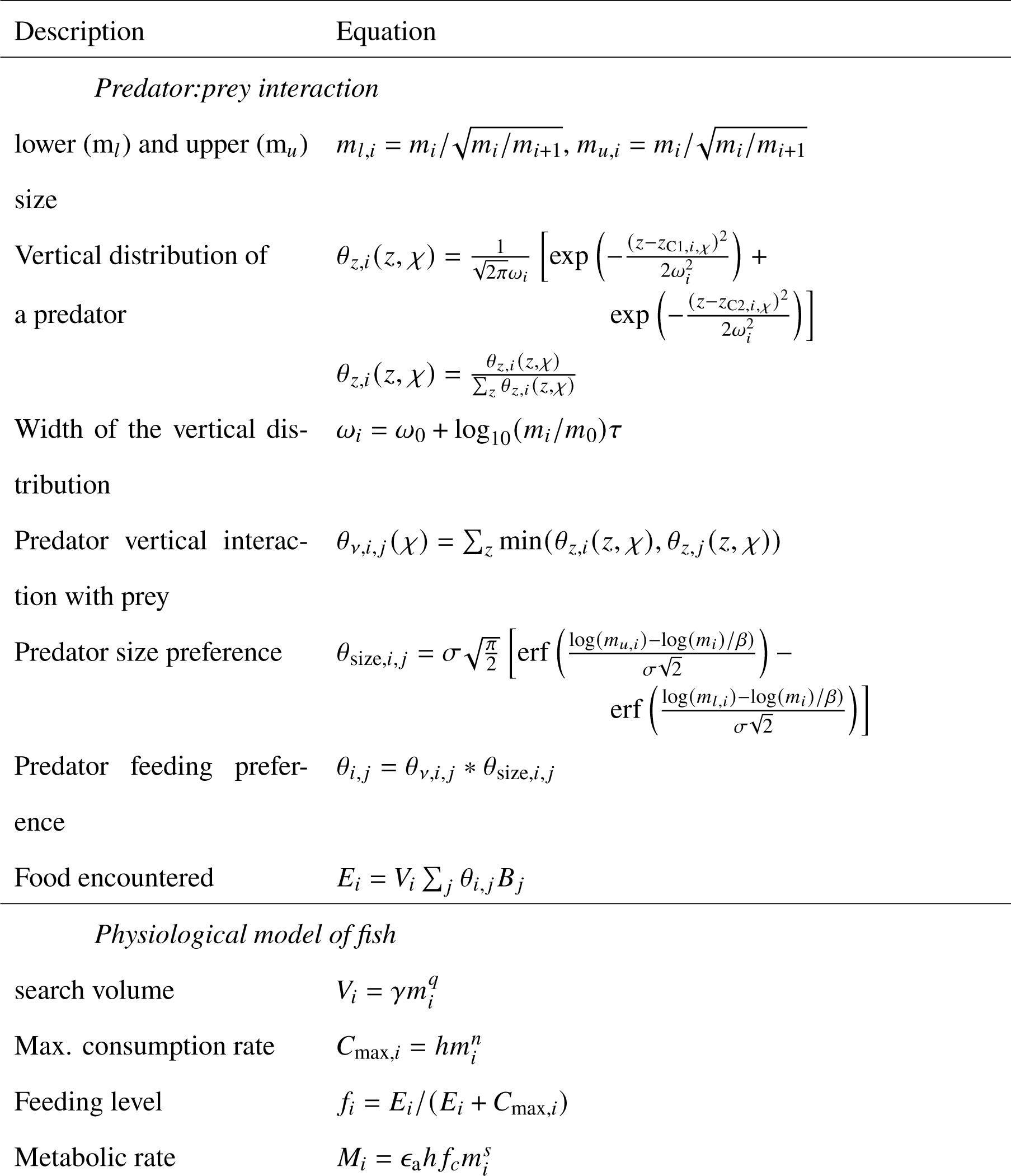
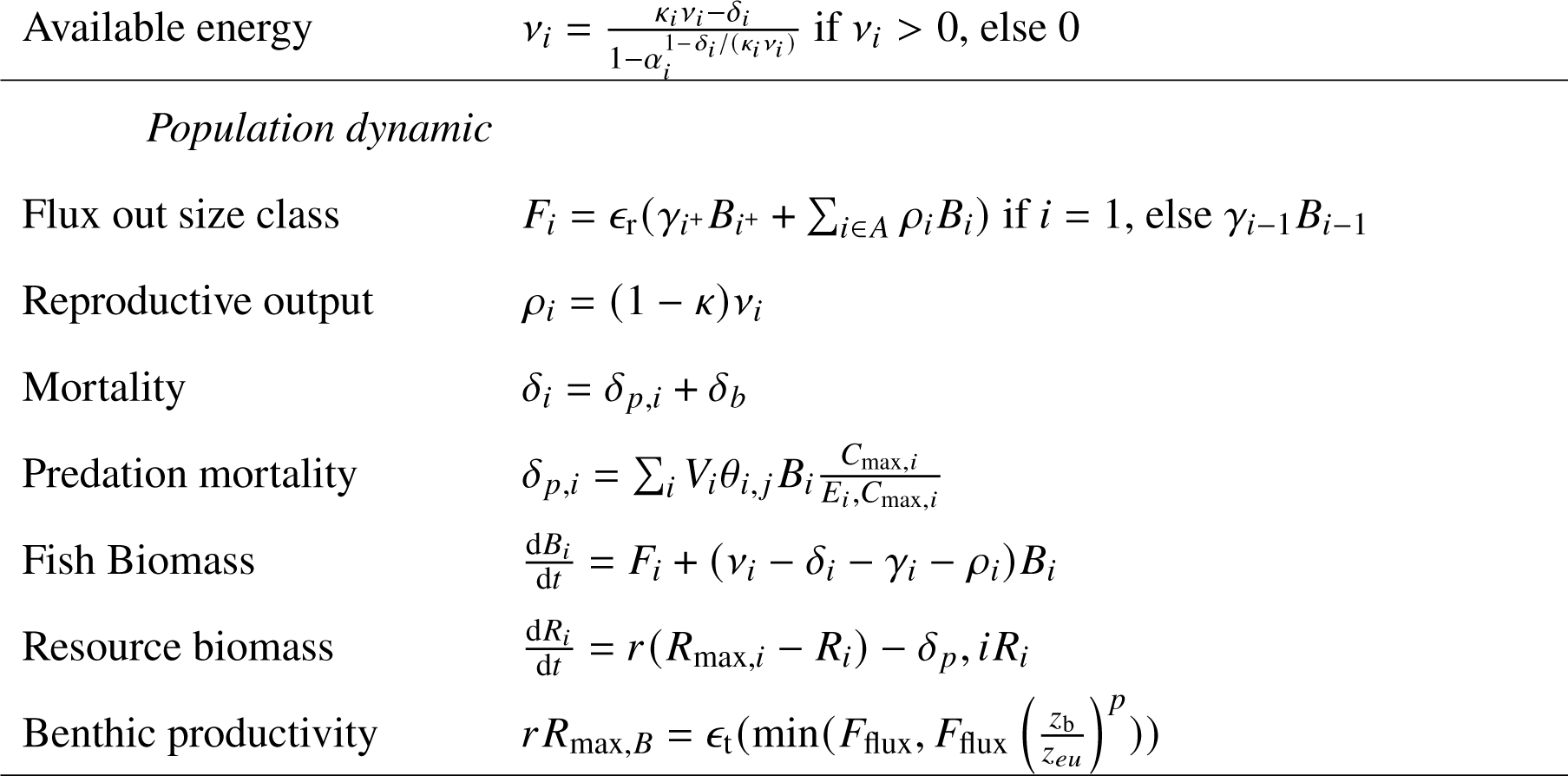
Main equation governing FEISTY.

**Table SB2:**
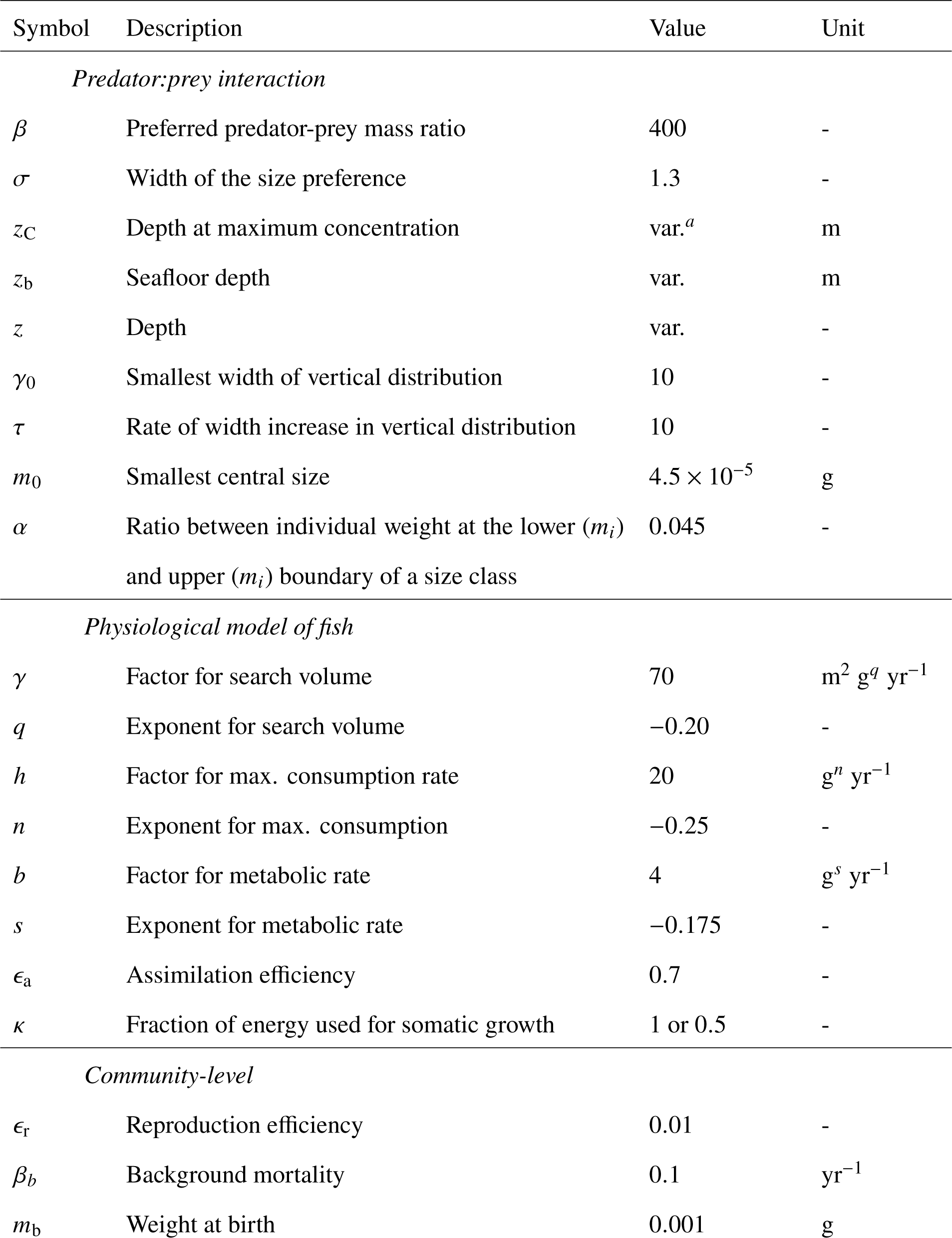
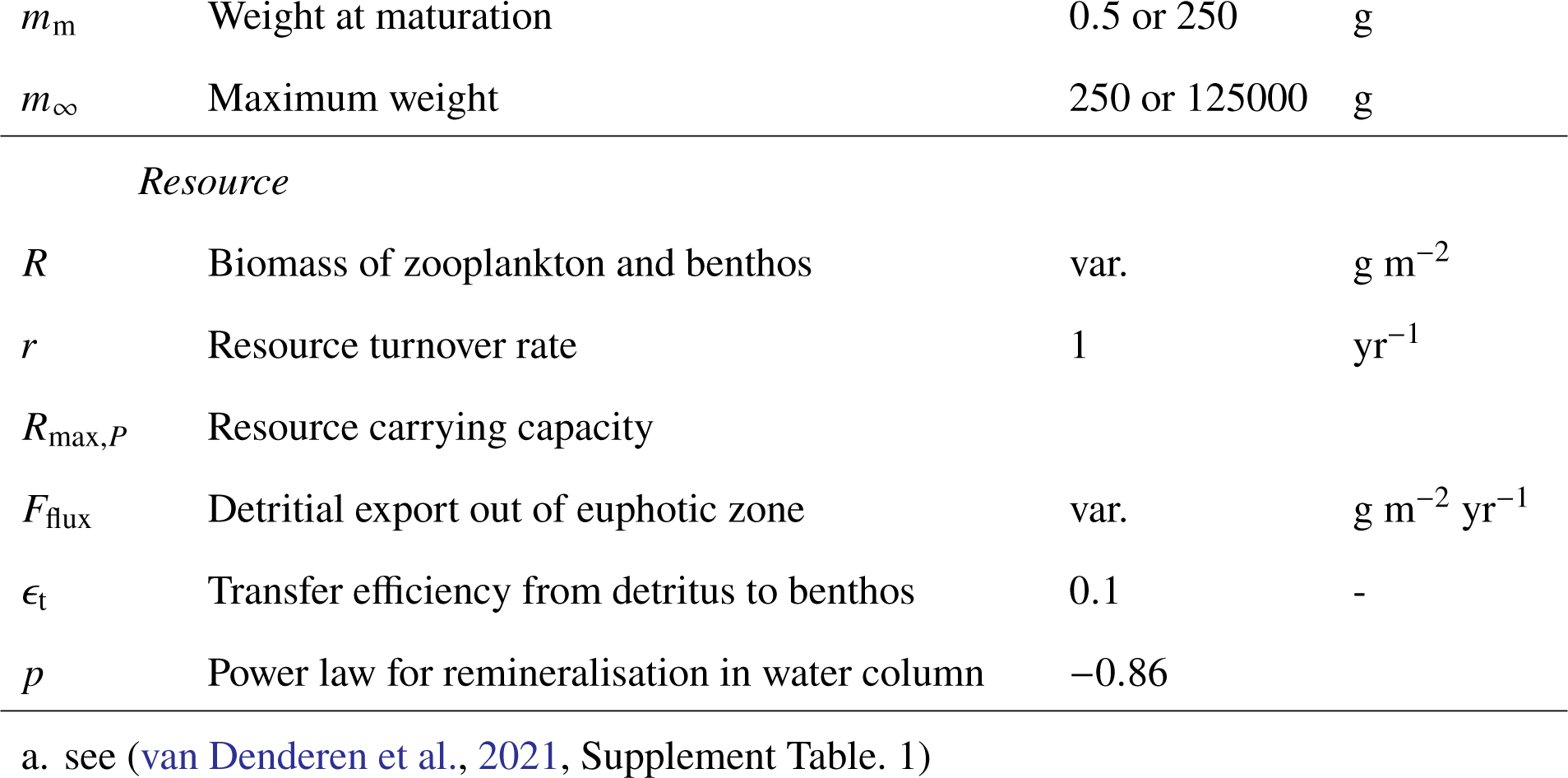
Parameters for fish in the FEISTY framework.

| Symbol | Description | Value | Unit |
| --- | --- | --- | --- |
| *Predator:prey interaction* |  |  |  |
| $\beta$ | Preferred predator-prey mass ratio | 400 | - |
| $\sigma$ | Width of the size preference | 1.3 | - |
| $z_C$ | Depth at maximum concentration | var.<sup>a</sup> | m |
| $z_b$ | Seafloor depth | var. | m |
| $z$ | Depth | var. | - |
| $\gamma_0$ | Smallest width of vertical distribution | 10 | - |
| $\tau$ | Rate of width increase in vertical distribution | 10 | - |
| $m_0$ | Smallest central size | $4.5 \times 10^{-5}$ | g |
| $\alpha$ | Ratio between individual weight at the lower ($m_i$) and upper ($m_i$) boundary of a size class | 0.045 | - |
| *Physiological model of fish* |  |  |  |
| $\gamma$ | Factor for search volume | 70 | $m^2 g^q yr^{-1}$ |
| $q$ | Exponent for search volume | -0.20 | - |
| $h$ | Factor for max. consumption rate | 20 | $g^n yr^{-1}$ |
| $n$ | Exponent for max. consumption | -0.25 | - |
| $b$ | Factor for metabolic rate | 4 | $g^s yr^{-1}$ |
| $s$ | Exponent for metabolic rate | -0.175 | - |
| $\epsilon_a$ | Assimilation efficiency | 0.7 | - |
| $\kappa$ | Fraction of energy used for somatic growth | 1 or 0.5 | - |
| *Community-level* |  |  |  |
| $\epsilon_r$ | Reproduction efficiency | 0.01 | - |
| $\beta_b$ | Background mortality | 0.1 | $yr^{-1}$ |
| $m_b$ | Weight at birth | 0.001 | g |
| $m_m$ | Weight at maturation | 0.5 or 250 | g |
| $m_\infty$ | Maximum weight | 250 or 125000 | g |
| *Resource* |  |  |  |
| $R$ | Biomass of zooplankton and benthos | var. | $\text{g m}^{-2}$ |
| $r$ | Resource turnover rate | 1 | $\text{yr}^{-1}$ |
| $R_{\max,P}$ | Resource carrying capacity | | |
| $F_{\text{flux}}$ | Detrital export out of euphotic zone | var. | $\text{g m}^{-2} \text{yr}^{-1}$ |
| $\epsilon_t$ | Transfer efficiency from detritus to benthos | 0.1 | - |
| $p$ | Power law for remineralisation in water column | -0.86 | |

a. see (van Denderen et al., 2021, Supplement Table. 1)

## Supplement C Cephalopods in Ecopath

We have reviewed data from seven Ecopath models that include cephalopods in the analysis (Table. SC1). For each model we classified the species or group in different functional group that correspond to the FEISTY functional group – Zooplankton, pelagic, demersals, and cephalopods. We reported the ratio of cephalopod biomass *B*_ceph_ vs pelagic fish *B*_pel_ estimated from Ecopath models, i.e., *B*_ceph_/(*B*_ceph_ + *B*_pel_). The zooplankton productivity (g m^−2^ yr^−1^) is calculated as product of the production rate over biomass (*P*/*Q* in yr^−1^) and the biomass of zooplankton (g m^−2^).

**Table SC1:** Ecopath models including cephalopods.

| ID | Maximum depth | Region | Ecosystem type | Reference |
| --- | --- | --- | --- | --- |
| 438 | 1000 | South East Alaska | Continental shelf | Guénette et al. (2006) |
| 443 | 2000 | West Scotland | Open ocean | Howell et al. (2009) |
| 448 | 150 | Irish Sea | Continental shelf | Lees and Mackinson (2007) |
| 485 | 500 | South Benguela | Upwelling | Shannon et al. (2003) |
| 733 | 600 | Celtic Sea-Biscay | Continental shelf | Bentorcha et al. (2017) |
| 735 | 200 | Celtic Sea | Continental shelf | Moullec et al. (2017) |
| 738 | 5000 | Azores | Open ocean | Morato et al. (2016) |

**Figure SC1:**
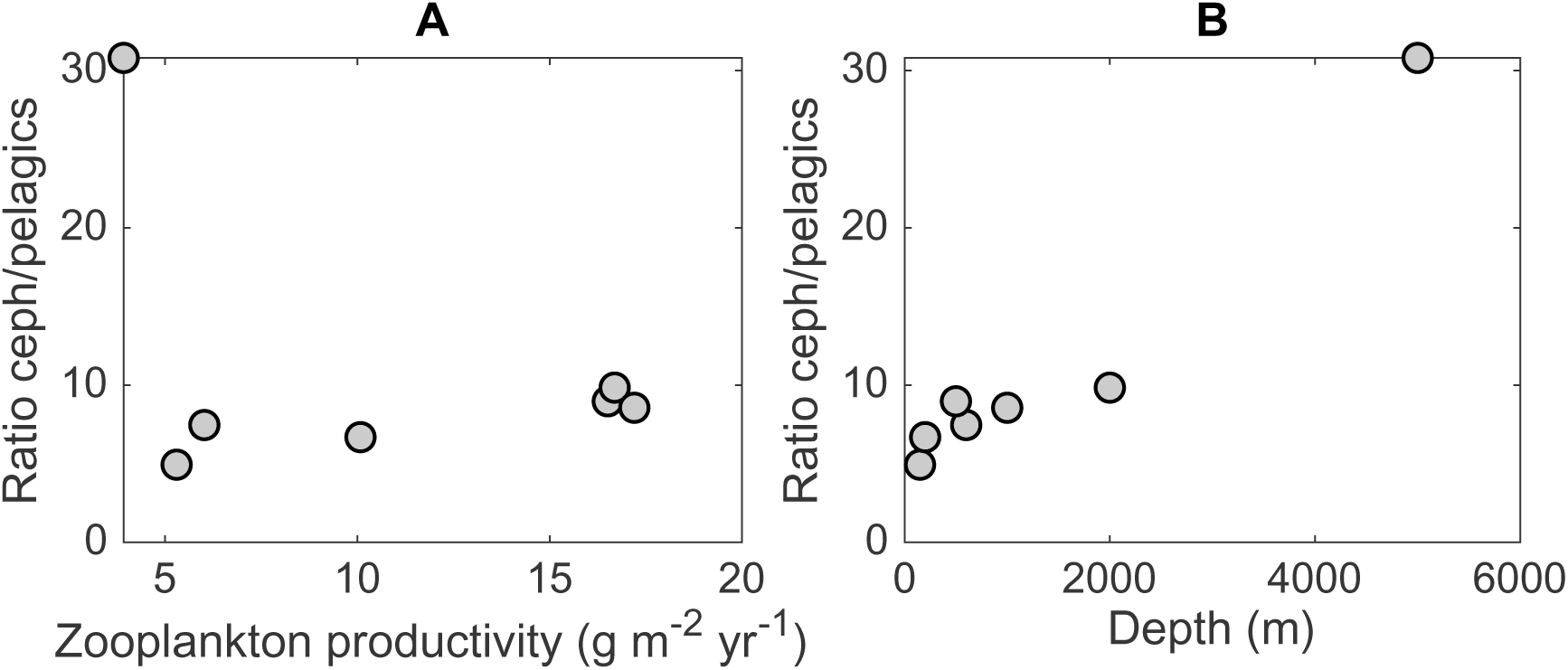
Ratio of cephalopods vs pelagic (in percent) in Ecopath models with secondary production (A) and depth (B)

## Supplement D Effect of predation by squid in FEISTY

The decline of total biomass due to the presence of squid (Fig.4) is not explain by competition in our model. Their presence results in similar feeding levels for demersal and small pelagic in shelf region (Fig.5E and F) and for small pelagic and large pelagic in open ocean (Fig.5G and H). We further show that increasing the intensity of predation from squid on pelagic groups – small and large – and demersal has a strong effect on other group biomass and total biomass in the system (Fig. SD1). We also show that increasing predation on pelagic in open ocean results in a change in composition of the system from a dominance of both large and small pelagic to a system dominated by epipelagic fish (Fig. SD1B).

**Figure SD1:**
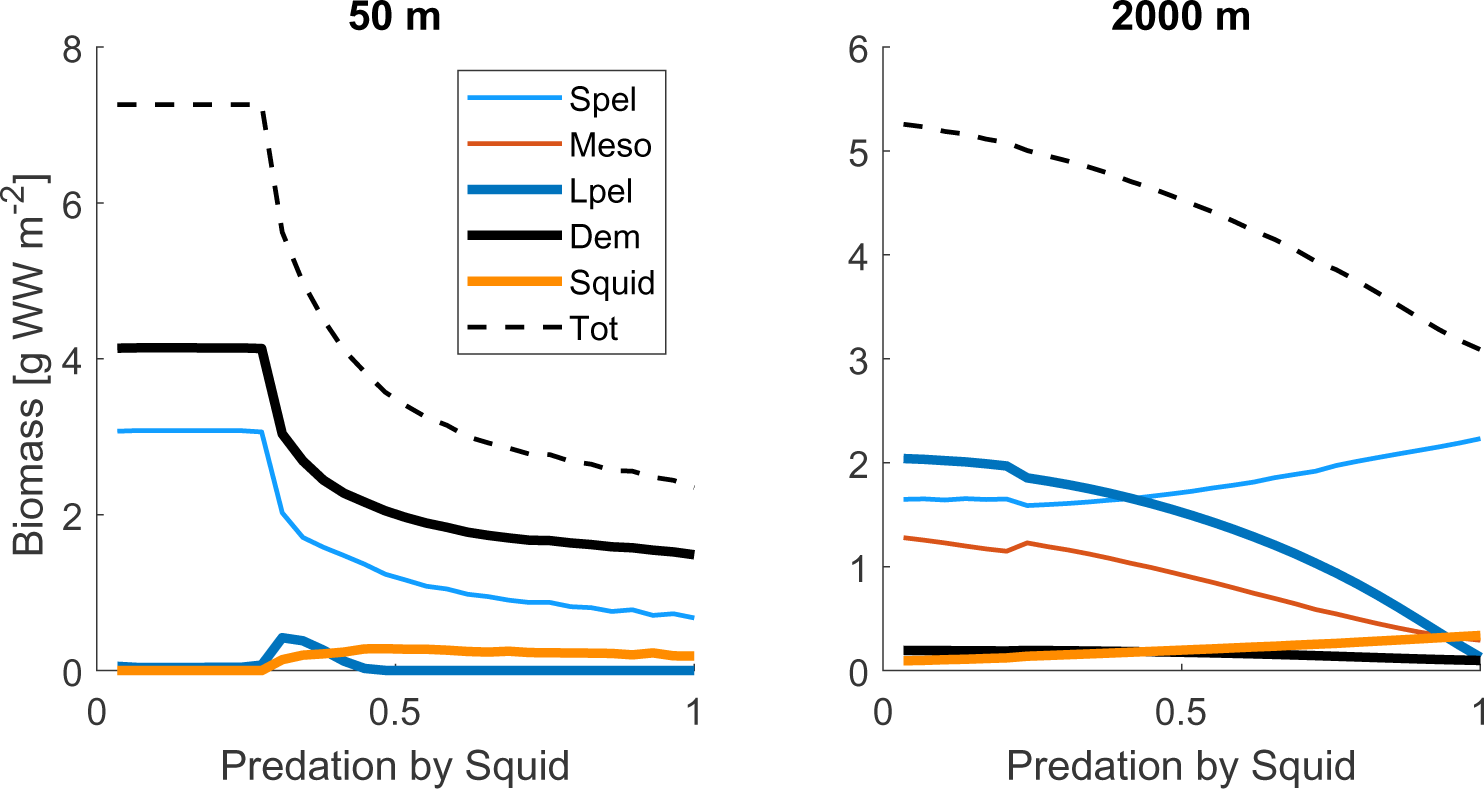
Effect of squid efficiency to prey on pelagic fish (epipelagic fish and large pelagic) and demersal. Efficiency of predation ranges from 0, no predation to 1, predation depends only on size- and vertical-overlap of squid and their prey. Simulation has been realised at 50, 2000 meters at a zooplankton productivity of 100 g m^−2^ yr^−1^.

## Supplement E Turnover rate of energy of mature organisms

**Figure SE1:**
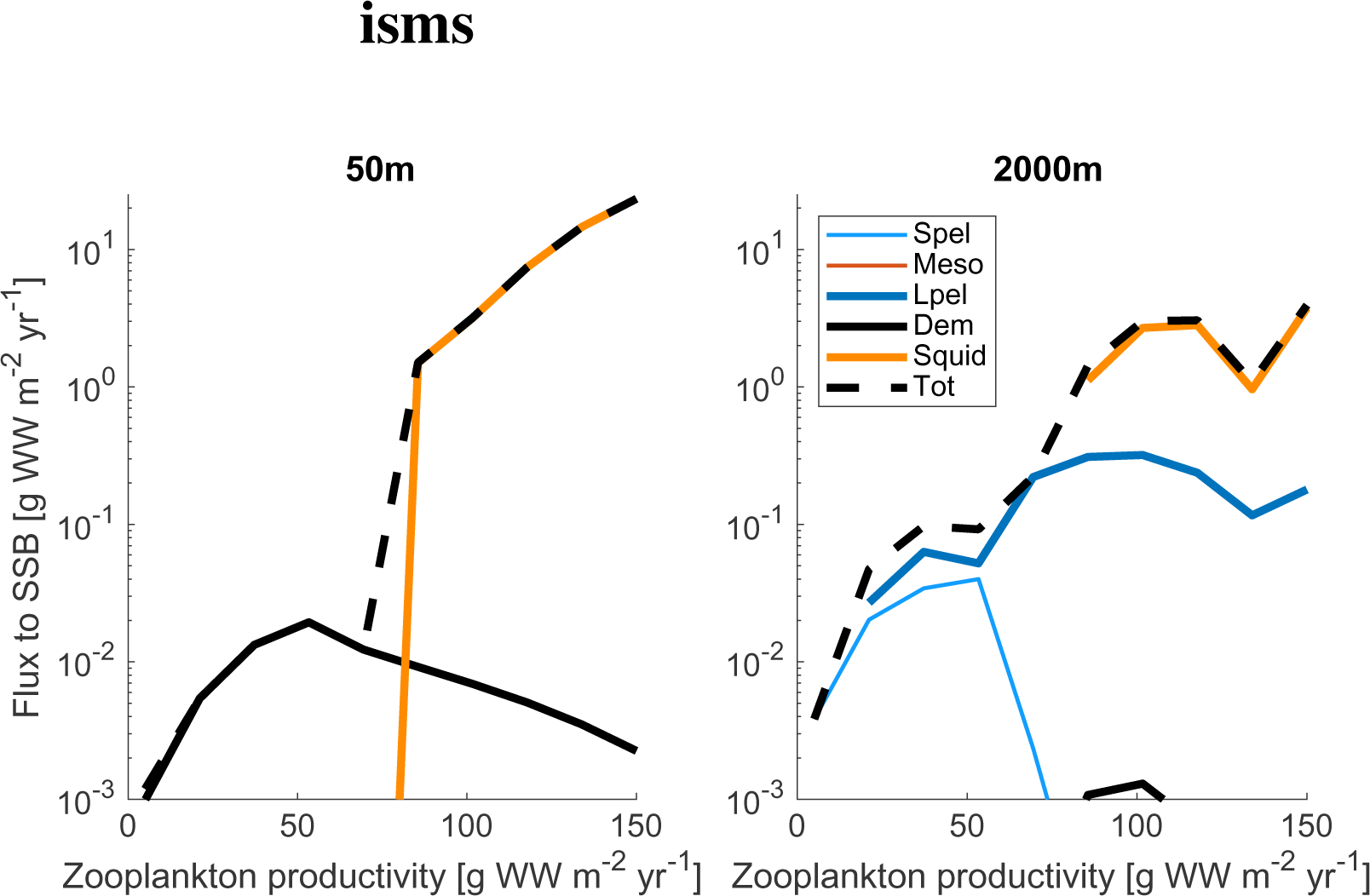
Mass flux to Spawning Stock Biomass (SSB) for a shelf (left panel) and an open ocean (right panel) region. The flux to SSB is calculated as the flux of a size class *i* weighted by the proportion of the matures in the following size class *i* 1 (i.e., *F*_SSB_ = *F*_*out*,*i*_1 − *k*_*i*+1_). Note that the *F*_SSB_ does not take into account the difference in accumulation of biomass of mature due to several spawning event in fish.

## Notes

https://github.com/RemyDenechere/The-role-of-squid-for-food-web-structure-and-community-level-metabolism/releases/tag/v0.beta

### Competing Interest Statement

The authors have declared no competing interest.

## References

Agus, B., M. Mereu, R. Cannas, A. Cau, E. Coluccia, M. C. Follesa, and D. Cuccu (2018). Age determination of loligo vulgaris and loligo forbesii using eye lens analysis. Zoomorphology 137(1), 63–70.

Andersen, K. H. (2019). Fish ecology, evolution, and exploitation: A New Theoretical Synthesis. Princeton University Press.

Andersen, K. H. and J. E. Beyer (2006). Asymptotic size determines species abundance in the marine size spectrum. The American Naturalist 168(1), 54–61. PMID: 16685635.

Arkhipkin, A. I. (2013). Squid as nutrient vectors linking southwest atlantic marine ecosystems. Deep Sea Research Part II: Topical Studies in Oceanography 95, 7–20.

Arkhipkin, A. I., P. G. Rodhouse, G. J. Pierce, W. Sauer, M. Sakai, L. Allcock, J. Arguelles, J. R. Bower, G. Castillo, L. Ceriola, et al. (2015). World squid fisheries. Reviews in Fisheries Science & Aquaculture 23(2), 92–252.

Arkhipkin, A. I. and N. V. Silvanovich (1997). Age, growth and maturation of the squid martialia hyadesi (cephalopoda, ommastrephidae) in the south-west atlantic. Antarctic Science 9(4), 373–380.

Bentorcha, A., D. Gascuel, and S. Guénette (2017). Using trophic models to assess the impact of fishing in the bay of biscay and the celtic sea. Aquatic Living Resources 30, 7.

Boyle, P. and S. Boletzky (1996). Cephalopod populations: definition and dynamics. Philosophical Transactions of the Royal Society of London. Series B: Biological Sciences 351(1343), 985–1002.

Boyle, P. and P. Rodhouse (2008). Cephalopods: ecology and fisheries. John Wiley & Sons.

Coll, M., J. Navarro, R. J. Olson, and V. Christensen (2013). Assessing the trophic position and ecological role of squids in marine ecosystems by means of food-web models. Deep Sea Research Part II: Topical Studies in Oceanography 95, 21–36.

de la Chesnais, T., E. A. Fulton, S. R. Tracey, and G. T. Pecl (2019). The ecological role of cephalopods and their representation in ecosystem models. Reviews in Fish Biology and Fisheries 29(2), 313–334.

De Roos, A. M., T. Schellekens, T. Van Kooten, K. Van De Wolfshaar, D. Claessen, and L. Persson (2008). Simplifying a physiologically structured population model to a stage-structured biomass model. Theoretical population biology 73(1), 47–62.

Denéchère, R., P. D. van Denderen, and K. H. Andersen (2022). Deriving population scaling rules from individual-level metabolism and life history traits. The American Naturalist 199(4), 564–575.

Doubleday, Z. A., T. A. Prowse, A. Arkhipkin, G. J. Pierce, J. Semmens, M. Steer, S. C. Leporati, S. Lourenço, A. Quetglas, W. Sauer, et al. (2016). Global proliferation of cephalopods. Current Biology 26(10), R406–R407.

Fang, Z., J. Li, K. Thompson, F. Hu, X. Chen, B. Liu, and Y. Chen (2016). Age, growth, and population structure of the red flying squid (ommastrephes bartramii) in the north pacific ocean, determined from beak microstructure. Fishery Bulletin 114(1).

Froese, R. and D. Pauly (2018). FishBase. *World wide web electronic publication.* www.fishbase.org.

Garibaldi, F. and M. Podestà (2014). Stomach contents of a sperm whale (physeter macrocephalus) stranded in italy (ligurian sea, north-western mediterranean). Journal of the Marine Biological Association of the United Kingdom 94(6), 1087–1091.

Goicochea-Vigo, C., E. Morales-Bojórquez, V. Y. Zepeda-Benitez, J. Á. Hidalgo-de-la Toba, H. Aguirre-Villaseñor, J. Mostacero-Koc, and D. Atoche-Suclupe (2019). Age and growth estimates of the jumbo flying squid (dosidicus gigas) off peru. Aquatic Living Resources 32, 7.

Guénette, S., S. J. Heymans, V. Christensen, and A. W. Trites (2006). Ecosystem models show combined effects of fishing, predation, competition, and ocean productivity on steller sea lions (eumetopias jubatus) in alaska. Canadian Journal of Fisheries and Aquatic Sciences 63(11), 2495–2517.

Hernández-Muñoz, A. T., C. Rodŕıguez-Jaramillo, A. Mejía-Rebollo, and C. A. Salinas-Zavala (2016). Reproductive strategy in jumbo squid dosidicus gigas (d’orbigny, 1835): A new perspective. Fisheries research 173, 145–150.

Hoving, H.-J. T. and B. Robison (2016). Deep-sea in situ observations of gonatid squid and their prey reveal high occurrence of cannibalism. Deep Sea Research Part I: Oceanographic Research Papers 116, 94–98.

Howell, K., S. Heymans, J. Gordon, M. Ayers, and E. Jones (2009). *DEEPFISH Project: Applying an ecosystem approach to the sustainable management of deep-water fisheries. Part 1: Development of an Ecopath with Ecosim model and Part 2: A new aproach to managing deep-water fisheries*. Number 259a, 259b in SAMS Internal reports. Scottish Association for Marine Science. Pages 116 Publisher SAMS Report no. 259.

Hunsicker, M. E., T. E. Essington, R. Watson, and U. R. Sumaila (2010). The contribution of cephalopods to global marine fisheries: can we have our squid and eat them too? Fish and fisheries 11(4), 421–438.

Hutchings, A. J., R. A. Myers, B. V. Garćıa, O. L. Lucifora, and A. Kuparinen (2012). Life-history correlates of extinction risk and recovery potential. Ecological Applications 22(4), 1061–1067.

Jackson, G. and M. Domeier (2003). The effects of an extraordinary el niño/la niña event on the size and growth of the squid loligo opalescens off southern california. Marine Biology 142(5), 925–935.

Kooijman, S. A. L. M. (2009). Dynamic Energy Budget Theory for Metabolic Organisation. New York: Cambridge University Press.

Lankford, T. E., J. M. Billerbeck, and D. O. Conover (2001). Evolution of intrinsic growth and energy acquisition rates. ii. trade-offs with vulnerability to predation in menidia menidia. Evolution 55(9), 1873–1881.

Laptikhovsky, V., A. Arkhipkin, M. R. Lipiński, U. Markaida, H. Murua, C. M. Nigmatullin, W. H. Sauer, and H.-J. T. Hoving (2019). Iteroparity or semelparity in the jumbo squid dosidicus gigas: a critical choice. Journal of Shellfish Research 38(2), 375–378.

Lees, K. and S. Mackinson (2007). An ecopath model of the irish sea: ecosystems properties and sensitivity analysis. Sci. Ser. Tech Rep..

MACY III, W. K. (1982). Feeding patterns of the long-finned squid, loligo pealei, in new england waters. The Biological Bulletin 162(1), 28–38.

Miyahara, K., T. Ota, T. Goto, and S. Gorie (2006). Age, growth and hatching season of the diamond squid thysanoteuthis rhombus estimated from statolith analysis and catch data in the western sea of japan. Fisheries Research 80(2-3), 211–220.

Morais, R. A. and D. R. Bellwood (2018). Global drivers of reef fish growth. Fish and Fisheries 19(5), 874–889.

Morato, T., E. Lemey, G. Menezes, C. K. Pham, J. Brito, A. Soszynski, T. J. Pitcher, and J. J. Heymans (2016). Food-web and ecosystem structure of the open-ocean and deep-sea environments of the azores, ne atlantic. Frontiers in Marine Science 3, 245.

Moullec, F., D. Gascuel, K. Bentorcha, S. Guénette, and M. Robert (2017). Trophic models: What do we learn about celtic sea and bay of biscay ecosystems? Journal of Marine Systems 172, 104–117.

Nabhitabhata, J. (1995, 01). Mass culture of cephalopods in thailand. World Aquaculture 26, 25–29.

Neuheimer, A. B., M. Hartvig, J. Heuschele, S. Hylander, T. Kiørboe, K. H. Olsson, J. Sainmont, and K. H. Andersen (2015). Adult and offspring size in the ocean over 17 orders of magnitude follows two life history strategies. Ecology 96(12), 3303–3311.

O’dor, R. and D. Webber (1986). The constraints on cephalopods: why squid aren’t fish. Canadian Journal of Zoology 64(8), 1591–1605.

Pecl, G. T. (2000). Comparative life history of tropical and temperate Sepioteuthis squids in Australian waters. Ph. D. thesis, James Cook University.

Pecl, G. T. and G. D. Jackson (2008). The potential impacts of climate change on inshore squid: biology, ecology and fisheries. Reviews in Fish Biology and Fisheries 18(4), 373–385.

Pérez-Palafox, X. A., E. Morales-Bojórquez, M. D. C. Rodŕıguez-Jaramillo, J. G. Díaz-Uribe, A. Hernández-Herrera, O. U. Rodŕıguez-Garćıa, and D. I. Arizmendi-Rodŕıguez (2019). Evidence of iteroparity in jumbo squid dosidicus gigas in the gulf of california, mexico. Journal of Shellfish Research 38(1), 149–162.

Persson, L., A. Van Leeuwen, and A. M. De Roos (2014). The ecological foundation for ecosystem-based management of fisheries: mechanistic linkages between the individual-, population-, and community-level dynamics. ICES Journal of Marine Science 71(8), 2268–2280.

Petrik, C. M., C. A. Stock, K. H. Andersen, P. D. van Denderen, and J. R. Watson (2019). Bottom-up drivers of global patterns of demersal, forage, and pelagic fishes. Progress in oceanography 176, 102124.

Phillips, K. L., G. D. Jackson, and P. D. Nichols (2001). Predation on myctophids by the squid moroteuthis ingens around macquarie and heard islands: stomach contents and fatty acid analyses. Marine Ecology Progress Series 215, 179–189.

Phillips, K. L., P. D. Nichols, and G. D. Jackson (2003). Size-related dietary changes observed in the squid moroteuthis ingens at the falkland islands: stomach contents and fatty-acid analyses. Polar Biology 26(7), 474–485.

Rodhouse, P., E. G. Dawe, and R. K. O’Dor (1998). Squid recruitment dynamics: the genus Illex as a model, the commercial Illex species and influence on variability, Volume 376. Food & Agriculture Org.

Rodhouse, P. and C. M. Nigmatullin (1996). Role as consumers. Philosophical Transactions of the Royal Society of London. Series B: Biological Sciences 351(1343), 1003–1022.

Rodhouse, P. G. (2005). Review of the state of world marine fishery resources: Fisheries technical paper. FAO Fisheries Tech. Paper 457, 175–187.

Rodhouse, P. G. and E. Hatfield (1990). Dynamics of growth and maturation in the cephalopod illex argentinus de castellanos, 1960 (teuthoidea: Ommastrephidae). Philosophical Transactions of the Royal Society of *London*. Series B: Biological Sciences 329(1254), 229–241.

Roper, C. F. and R. E. Young (1975). Vertical distribution of pelagic cephalopods.

Rosa, R., C. Yamashiro, U. Markaida, P. Rodhouse, C. M. Waluda, C. A. Salinas-Zavala, F. Keyl, R. O’Dor, J. S. Stewart, and W. F. Gilly (2013). Dosidicus gigas, humboldt squid.

Ryther, J. H. (1969). Photosynthesis and fish production in the sea: The production of organic matter and its conversion to higher forms of life vary throughout the world ocean. Science 166(3901), 72–76.

Schramski, J. R., A. I. Dell, J. M. Grady, R. M. Sibly, and J. H. Brown (2015). Metabolic theory predicts whole-ecosystem properties. Proceedings of the National Academy of Sciences 112(8), 2617–2622.

Seibel, B. A. (2007). On the depth and scale of metabolic rate variation: scaling of oxygen consumption rates and enzymatic activity in the class cephalopoda (mollusca). Journal of Experimental Biology 210(1), 1–11.

Shannon, L. J., C. L. Moloney, A. Jarre, and J. G. Field (2003). Trophic flows in the southern benguela during the 1980s and 1990s. Journal of Marine Systems 39(1-2), 83–116.

Smale, M. (1996). Cephalopods as prey. iv. fishes. Philosophical Transactions of the Royal Society of London. Series B: Biological Sciences 351(1343), 1067–1081.

Stawitz, C. C. and T. E. Essington (2019). Somatic growth contributes to population variation in marine fishes. Journal of Animal Ecology 88(2), 315–329.

Stock, C. A., J. G. John, R. R. Rykaczewski, R. G. Asch, W. W. Cheung, J. P. Dunne, K. D. Friedland, V. W. Lam, J. L. Sarmiento, and R. A. Watson (2017). Reconciling fisheries catch and ocean productivity. Proceedings of the National Academy of Sciences 114(8), E1441–E1449.

Summers, W. C. (1971). Age and growth of loligo pealei, a population study of the common atlantic coast squid. The Biological Bulletin 141(1), 189–201.

van Denderen, D., H. Gislason, J. van den Heuvel, and K. H. Andersen (2020). Global analysis of fish growth rates shows weaker responses to temperature than metabolic predictions. Global Ecology and Biogeography 29(12), 2203–2213.

van Denderen, P. D., C. M. Petrik, C. A. Stock, and K. H. Andersen (2021). Emergent global biogeography of marine fish food webs. Global Ecology and Biogeography 30(9), 1822–1834.

Villanueva, R., E. A. Vidal, F. Fernández-Á lvarez, and J. Nabhitabhata (2016). Early mode of life and hatchling size in cephalopod molluscs: Influence on the species distributional ranges. PLoS ONE 11(11), 1–27.

Vovk, A. (1985). Feeding spectrum of longflin squid (loligo pealei) in the northwest atlantic and its position in the ecosystem. NAFO Sci. Coun. Studies 3, 33–38.

William, G., C. Baxter, B. Block, A. Boustany, L. Zeidberg, K. Reisenbichler, B. Robison, G. Bazzino Ferreri, and C. A. Zavala (2006, 11). Vertical and horizontal migrations by squid dosidicus gigas revealed by electronic tagging. Marine Ecology Progress Series 324, 1–17.

